# Mapping host-microbe transcriptional interactions by dual perturb-seq

**DOI:** 10.1101/2023.04.21.537779

**Authors:** Simon Butterworth, Kristina Kordova, Sambamurthy Chandrasekaran, Kaitlin K. Thomas, Francesca Torelli, Eloise J. Lockyer, Amelia Edwards, Robert Goldstone, Anita A. Koshy, Moritz Treeck

## Abstract

Intracellular pathogens and other endosymbionts reprogram host cell transcription to suppress immune responses and recalibrate biosynthetic pathways. This reprogramming is critical in determining the outcome of infection or colonisation. Here, we combine pooled CRISPR knockout screening with dual host–microbe single-cell RNA-sequencing to identify the molecular mediators of these transcriptional interactions, a method we term dual perturb-seq. Applying dual perturb-seq to the intracellular pathogen *Toxoplasma gondii*, we are able to identify previously uncharacterised effector proteins and directly infer their function from the transcriptomic data. We show that *Tg*GRA59 contributes to the export of other effector proteins from the parasite into the host cell and identify a novel effector, *Tg*SOS1, that is necessary for sustained host STAT6 signalling and thereby contributes to parasite immune evasion and persistence. Together, this work demonstrates a novel tool that can be broadly adapted to interrogate host-microbe transcriptional interactions and reveal mechanisms of infection and immune evasion.

## INTRODUCTION

Endosymbiosis, a phenomenon in which one organism resides within the cells of another, is widespread across all domains of life and spans a continuum from mutually beneficial to pathogenic^1^. Establishment of endosymbiosis commonly involves transcriptional reprogramming of the host cell to downregulate immune responses and maintain a favourable environment for the endosymbiont^2^. Modulation of the host cell is often mediated by effector proteins secreted by the endosymbiont that directly interface with host signalling pathways^3, 4^.

The single-celled eukaryotic parasite *Toxoplasma gondii* is outstanding in both the number of identified effector proteins that target host cell transcription and breadth of this reprogramming. While bacterial effector proteins are exported from the cytosol through molecular secretion systems, *T. gondii* delivers effector proteins into the host cell from two types of secretory organelles, rhoptries and dense granules. Rhoptry proteins are injected directly into the host cytoplasm during parasite invasion of a host cell. Dense granule proteins are secreted into the parasitophorous vacuole in which *T. gondii* resides following host cell invasion. Some secreted dense granule proteins are then transported into the host cell via a secondary export mechanism^5^. More than 200 proteins have been localised to the rhoptries and dense granules of *T. gondii*, of which the vast majority are uncharacterised^6^. Most of these proteins represent novel orthologue groups found only in the Apicomplexa phylum of intracellular parasites, or sub-groups thereof. Many do not contain recognisable conserved functional domains that would inform on function^6^. Linking *T. gondii*-induced transcriptional responses in the host cell to specific effector proteins is thus highly challenging. Moreover, existing competitive growth-based pooled knockout screens in *T. gondii* have failed to identify effector proteins known to target host cell transcription, despite their critical roles in parasite virulence, indicating that a novel methodology is needed^7–10^.

Here, we adapt a method for pooled CRISPR-Cas9 knockout screening with a single-cell transcriptome readout to measure the impact of gene knockouts in an intracellular pathogen on the infected host cell, which we term dual perturb-seq. We demonstrate this method in *T. gondii* using a plasmid vector that enables the direct capture of both mRNAs and sgRNAs. Single-cell transcriptomic profiles of host cells infected with genetically perturbed parasites accurately recapitulate bulk RNA-seq phenotypes of established effector proteins. We screen a library of >1000 sgRNAs targeting 256 parasite genes encoding secreted rhoptry and dense granule proteins to gain the first system-level view of host cell reprogramming by the *T. gondii* secretome. This screen identifies all previously known effector proteins that modulate host cell transcription and additional putative effectors. We validate the dual perturb-seq phenotypes for two such effectors, showing that GRA59 contributes to export of dense granule effectors into the host cell, and that the previously unknown effector SOS1 is essential for sustained STAT6 signalling and M2 polarisation of infected cells.

## RESULTS

### Optimisation of a direct-capture perturb-seq vector for *Toxoplasma gondii*

Single cell CRISPR screening methods require identification of the sgRNA protospacer sequence to assign an identity to each cell. In direct-capture perturb-seq, the insertion of a “capture sequence” allows for direct sequencing of the sgRNA transcript in addition to the polyadenylated mRNA transcriptome following droplet-based cell partitioning and molecular barcoding^11^.

To enable direct-capture perturb-seq in *T. gondii*, we modified our existing Cas9-sgRNA plasmid vector^7^ by introducing one of two validated sgRNA-capture sequence configurations **(Figure 1A)**. Following transfection of these vectors into *T. gondii* by electroporation, we did not find any significant difference in knockout efficiency of the non-essential *T. gondii* UPRT gene for either sgRNA compared to the unmodified sgRNA, as measured by knockout-induced resistance to the toxic nucleoside analogue 5-fluorodeoxyuridine^12^ **(Figure 1B)**. As sgRNA-CS2 had a higher mean knockout efficiency than sgRNA-CS1, we used this configuration for our initial validation experiment.

**Figure 1.**
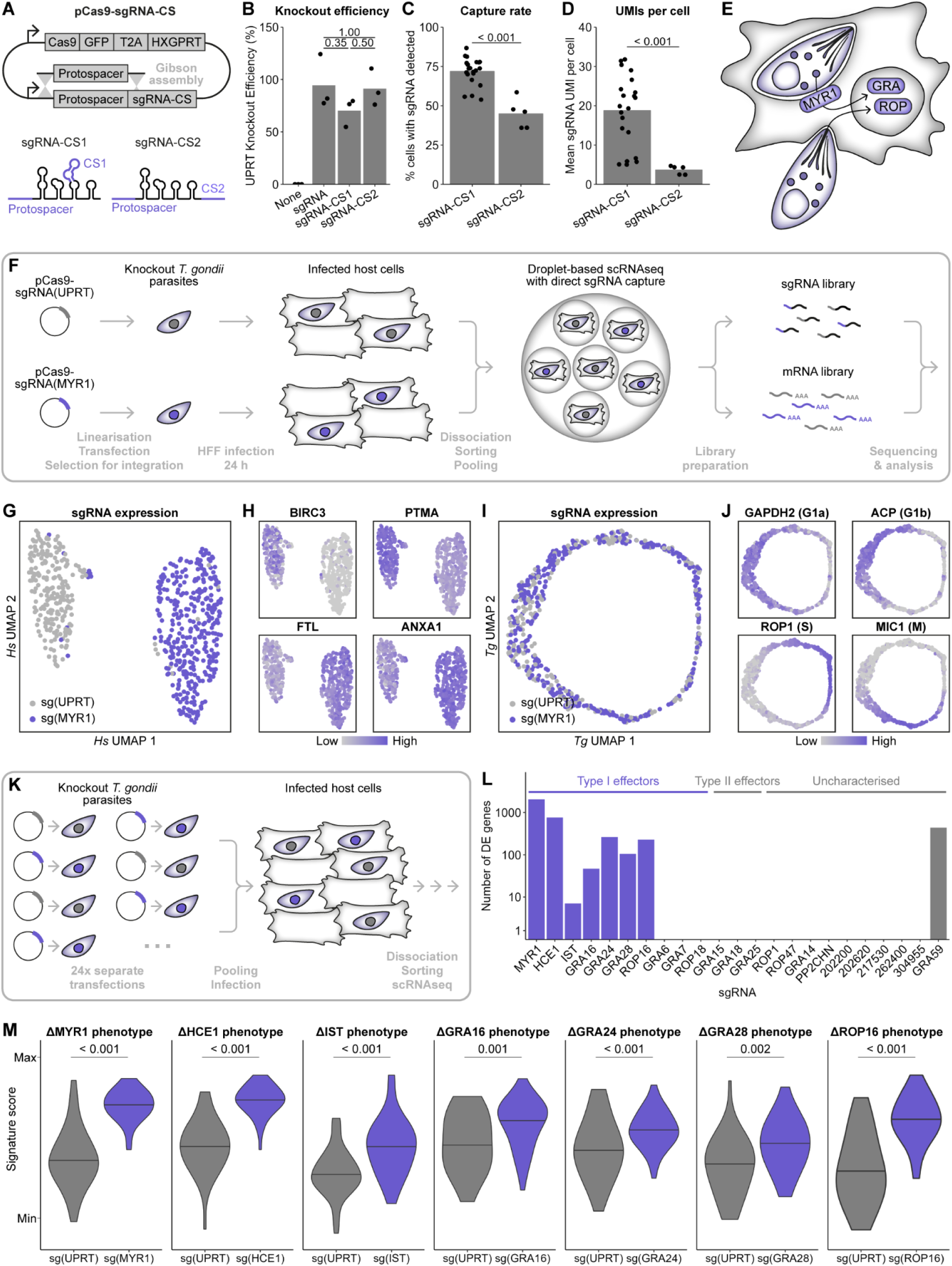
Dual perturb-seq transcriptional profiles recapitulate phenotypes of known *T. gondii* effectors. **A.** Schematic of perturb-seq plasmid vector and capture sequence sgRNAs. **B.** Knockout efficiency of perturb-seq vectors. Loss of function of the *T. gondii* UPRT gene was measured by plaque assay in the presence/absence of 5-fluorodeoxyuridine. Differences were tested by two-sided *t-*test with Bonferroni correction. **C.** Percentage of infected cells with a detectable sgRNA in all dual perturb-seq samples. Difference tested by two-sided *t-*test. **D.** Mean number of sgRNA UMIs in cells with a detectable sgRNA in all dual perturb-seq samples. Difference tested by two-sided *t-*test. **E.** Schematic of *T. gondii* effector protein export into the host cell. **F.** Experimental strategy for dual perturb-seq experiment with two sgRNAs, targeting UPRT and MYR1. **G.** UMAP of single cell transcriptomes based on host cell gene expression. **H.** Expression of two most significantly up-and down-regulated host cell marker genes for sg(MYR1)-expressing cells. **I.** UMAP of single cell transcriptomes based on *T. gondii* gene expression. **J.** Expression of *T. gondii* cell cycle marker genes. **K.** Experimental strategy for dual perturb-seq experiment with 24 sgRNAs. **L.** Number of significantly differentially expressed genes (p < 0.05) for each sgRNA compared to sg(UPRT). Differences tested by two-sided Wilcoxon rank-sum test with Benjamini-Hochberg adjustment. See also **Supplementary Data 2**. **M.** Scoring of single cell transcriptomes based on marker genes of established effector proteins using VISION. Differences tested by two-sided Wilcoxon rank-sum test.

However, across all experiments we completed as part of this work, we found that the capture rate of sgRNA-CS1 was significantly better than sgRNA-CS2 in terms of the percentage of cells in which any sgRNA is detected **(Figure 1C)** and the number of sgRNA UMIs detected **(Figure 1D)**. Therefore, in later experiments we used sgRNA-CS1 to maximise the number of recovered cells with an identifiable knockout.

### Dual perturb-seq transcriptional profiles recapitulate phenotypes of known *T. gondii* effectors

As perturb-seq has not been previously attempted in a non-mammalian system, we carried out two experiments to validate the dual perturb-seq transcriptional profiles of host cells infected with *T. gondii* mutants. First, we targeted the *T. gondii* UPRT gene, as a negative control, and the MYR1 gene, as a positive control. MYR1 is essential for export of dense granule effectors into the host cell, and it has been shown that knockout of MYR1 abrogates the majority of host cell transcriptional changes induced by *T. gondii*^13, 14^ **(Figure 1E)**. *T. gondii* RHΔHXGPRT parasites were transfected with perturb-seq plasmid vectors targeting either UPRT or MYR1, then used to infect human foreskin fibroblasts (HFFs) for 24 h at a multiplicity of infection (MOI) of 0.1 **(Figure 1F)**. Infected cells were sorted based on Cas9-GFP expression in the intracellular parasites and pooled for scRNA-seq using the 10x Genomics Chromium platform.

Sequencing reads were aligned to a dual *H. sapiens* and *T. gondii* reference and the cell barcodes were filtered to retain only infected cells in which a single sgRNA species was detected. **(Figure S1 and S2A)**. Inspecting these single cell transcriptomes by uniform manifold approximation projection (UMAP) using only the host cell gene expression, we observed that cells expressing the MYR1-targeting sgRNA formed a distinct cluster from those expressing the UPRT-targeting sgRNA, indicating that knockout of these genes in the intracellular parasite results in differing transcriptional profiles in the host cell **(Figure 1G)**. More than 5,000 host cell genes were significantly differentially expressed between cells expressing the MYR1-targeting and UPRT-targeting sgRNAs, including many previously shown to be dependent on MYR1^14^ **(Figure 1H and S2B, Supplementary Data 1A)**.

In each infected host cell, we were also able to resolve the transcriptome of the synchronous intracellular parasites. Projecting the cells by UMAP using the *T. gondii* gene expression, we found that the transcriptomes resolved in a circular structure that recapitulated the parasite cell cycle **(Figure 1I and 1J)**. Very few *T. gondii* genes were differentially expressed between cells expressing the MYR1-targeting and UPRT-targeting sgRNAs, indicating that knockout of these genes does not induce broad changes in the parasite transcriptome **(Figure S2C, Supplementary Data 1B)**.

To further validate this dual perturb-seq approach for identifying *T. gondii* effector proteins, we generated a panel of 22 vectors targeting 12 effector proteins that have been shown to impact host cell transcription and 10 rhoptry and dense granule proteins for which the host cell transcriptome has not been characterised. We transfected *T. gondii* RHΔHXGPRT separately with the 22 new vectors and the MYR1-and UPRT-targeting vectors and pooled the resulting transfectants into a single mixed population. This parasite pool was used to infect HFFs for 24 h, following which infected cells were enriched by fluorescence-activated cell sorting (FACS) for scRNA-seq as before **(Figure 1K)**. We recovered between 10 and 72 single cell transcriptomes for each sgRNA **(Figure S1D)**. Cells expressing different sgRNAs did not segregate into distinct clusters in a UMAP of host gene expression, although different sgRNAs were enriched in different regions of a large main cluster **(Figure S2E and S2F)**.

We identified differentially expressed host genes for seven out of ten effector proteins that have been validated in the highly virulent type I *T. gondii* RH strain used in this experiment: MYR1^14^, HCE1/TEEGR^15, 16^, IST^17, 18^, GRA16^19^, GRA24^20^, GRA28^21, 22^, and ROP16^23^ **(Figure 1L, Supplementary Data 2)**. We did not find any significantly differentially expressed genes for the effectors GRA6^24^, GRA7^25^ or ROP18^26^. These were among the genes for which we recovered the fewest single cell transcriptomes and so have less statistical power; however, there is also less evidence that these effectors induce changes in the host cell transcriptome during a physiological infection. There were no differentially expressed host genes for the type II *T. gondii* strain-specific effector GRA15^27^, as expected, nor for GRA18^28^ or GRA25^29^, suggesting that these are also type II-specific effectors. Of the uncharacterised genes included in this experiment, we identified differentially expressed host genes only for GRA59, which we investigate further later in this study.

Bulk transcriptomic datasets of *T. gondii* knockout strain-infected versus wild-type-infected host cells are available for the seven type I effectors for which we can identify differentially expressed genes, but not for GRA6, GRA7, or ROP18 (see **Methods**). To determine whether the host cell phenotype derived from this dual perturb-seq experiment concur with these datasets, we scored the single cell transcriptomes for the expression of marker genes identified in these prior studies using the VISION package^30^. Cells expressing the sgRNA targeting each of these seven effectors (MYR1, HCE1, IST, GRA16, GRA24, GRA28, and ROP16) had significantly higher scores for the cognate bulk transcriptome signatures compared to sg(UPRT)-expressing cells **(Figure 1M)**. Thus, we show that dual perturb-seq is able to deconvolve the phenotypes of host cells infected with a pool of different *T. gondii* knockouts and that the resulting transcriptomic profiles concur with all available prior datasets.

### Identification of *T. gondii* genes that alter host cell transcription through a dual perturb-seq screen

Having validated the dual perturb-seq workflow, we set up a knockout screen to identify all *T. gondii* effectors that target host cell transcription. We selected 256 rhoptry and dense granule localised proteins comprising the *T. gondii* effectome, based on spatial proteomics data^6^ and community annotation^31^. Five protospacer sequences targeting each of these genes were selected from our arrayed library^7^ and incorporated into the *T. gondii* perturb-seq vector by pooled Gibson assembly **(Supplementary Data 3)**. The plasmid pool was transfected into *T. gondii* RHΔHXGPRT, and the resulting pool of knockout parasites was used to infect HFFs **(Figure 2A)**. We carried out the dual perturb-seq screen in both unstimulated and interferon-gamma (IFNγ)-stimulated HFFs to identify effector proteins that modulate the expression of interferon-stimulated genes^17, 18, 32^.

**Figure 2:**
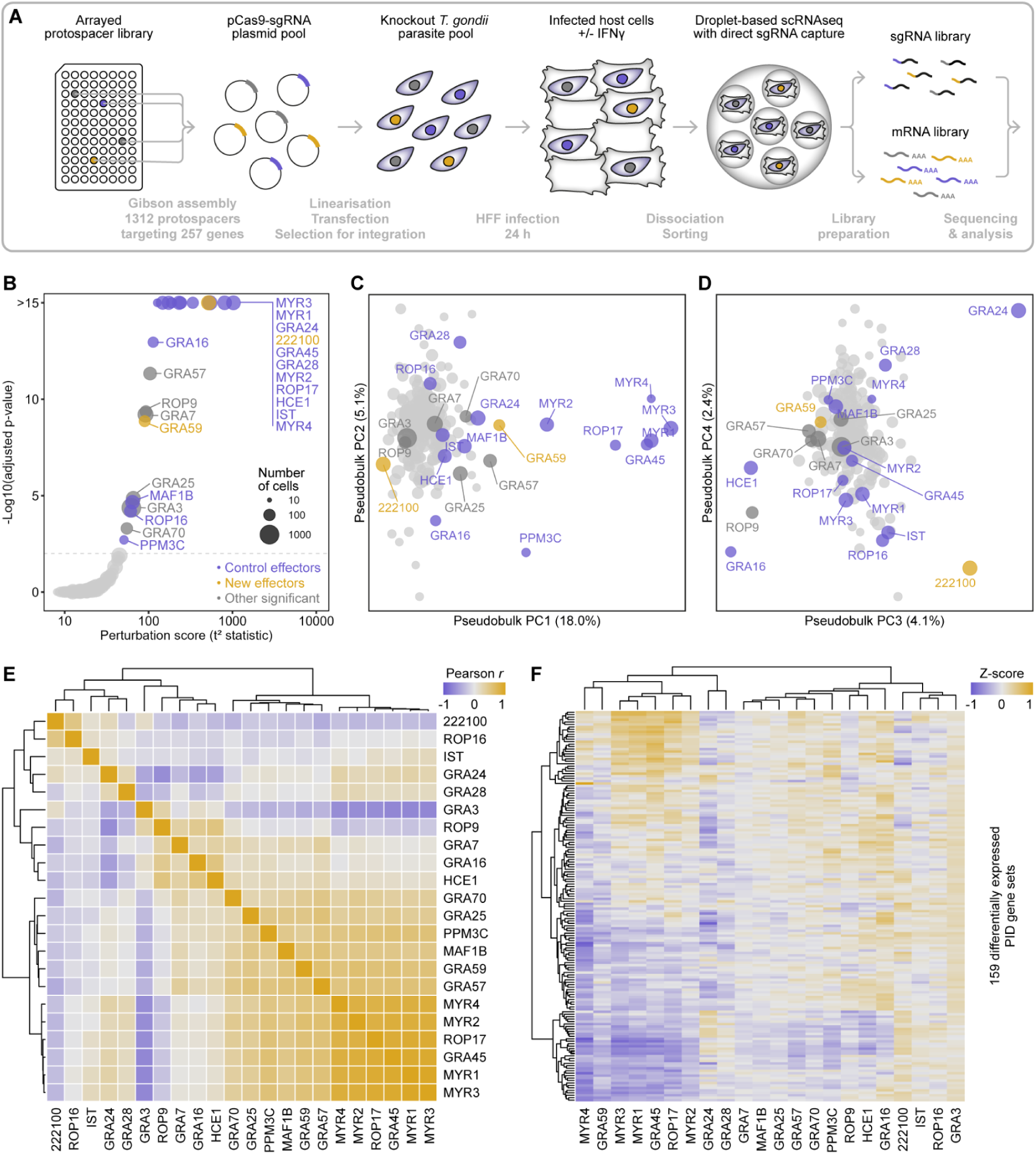
A dual-perturb-seq screen identifies *T. gondii* effectors that target host cell transcription. **A.** Schematic of dual perturb-seq screen. **B.** Perturbation of host cell transcriptomes by *T. gondii* effectors, measured by Hotelling’s *t*^2^-test on PCA embeddings of single cell transcriptomes with Benjamini Hochberg adjustment. See also **Supplementary Data 4**. **C.** PCA embeddings of pseudo-bulk transcriptomes. **D.** PCA embeddings of pseudo-bulk transcriptomes. **E.** Pearson correlations between pseudo-bulk transcriptomes of significant effectors. **F.** Average VISION signature scores of Pathway Interaction Database gene sets that are significantly differentially regulated by at least one significant effector (p < 0.01, two-sided Wilcoxon rank-sum test with Benjamini-Hochberg adjustment). See also **Figure S4** and **Supplementary Data 7**.

We recovered a total of 25,185 single-cell transcriptomes of HFFs infected with the pool of *T. gondii* mutants (15,920 unstimulated and 9,265 IFNγ-stimulated) **(Figure S3A and S3B)**. Across all the single cell transcriptomes we identified 1,119 sgRNAs with a median of 11 transcriptomes per sgRNA **(Figure S3E)**. All of the 256 *T. gondii* target genes were represented by at least one sgRNA; aggregating all sgRNAs targeting a given gene resulted in a median coverage of 71 transcriptomes per gene **(Figure S3F)**. Most target genes were represented by five different sgRNAs, and 93% of target genes were represented by at least three different sgRNAs **(Figure S3G)**. Both at the individual sgRNA level, and when aggregated by target gene, the number of single-cell transcriptomes recovered was directly proportional to sgRNA representation in the perturb-seq plasmid pool as assessed by bulk sequencing **(Supplementary Data 3)**, although sgRNAs targeting genes with strongly negative fitness phenotypes ^33^ tended to be less well represented **(Figure S3H and S3I)**.

In order to identify *T. gondii* gene knockouts that perturb the host cell transcriptome using a single metric, we analysed the single cell transcriptomes by principal component analysis (PCA) and tested for each target gene whether the distribution of single cell transcriptomes in PCA space differed from the background distribution using Hotelling’s *t*^2^-test, a multivariate generalisation of Student’s *t*-test^34^ **(Figure S3J)**. Combining the data from both unstimulated and IFNγ-stimulated cells, we identified 22 *T. gondii* genes which significantly alter the host cell transcriptome, including all 14 controls **(Figure 2B, Supplementary Data 4)**. We also analysed the data from unstimulated and IFNγ-stimulated cells separately, but did not identify any significant genes specific to the IFNγ-stimulated condition apart from the controls IST^17, 18^ and PPM3C^35^ **(Figure S3K and S3L, Supplementary Data 4)**. We identified significant perturbations for both GRA7 and GRA25, despite not identifying any DEGs for these knockouts in our earlier pilot experiment. Potentially, the higher coverage in this main screen in terms of number of single cell transcriptomes, and the use of multiple sgRNAs for each target, allows us to identify more subtle phenotypes. Most interestingly, we identified six genes which have not previously been implicated in reprogramming host transcription: TGGT1_222100, GRA57, ROP9, GRA59, GRA3, and GRA70.

We then applied Hotelling’s *t*^2^-test to single-cell PCA embeddings using the *T. gondii* transcriptome to test whether deletion of any parasite effector alters parasite gene transcription. We only found a significant perturbation for GRA3 **(Figure S4B, Supplementary Data 5)**. Inspection of genes differentially regulated in sg(GRA3)-expressing cells revealed that this perturbation was primarily driven by downregulation of GRA3 itself, with no other log2 fold-changes greater than 0.11 **(Figure S4C, Supplementary Data 6)**. Thus, knockout of effector proteins does not appear to affect the *T. gondii* transcriptome.

### Dual perturb-seq data reveals putative effector protein functions

To further investigate these putative effector proteins, we generated pseudo-bulk transcriptomes by averaging the Z-transformed normalised UMI counts for all target genes represented by at least 30 cells, corresponding to the 25^th^ percentile of target genes. Using PCA, we found that the majority of variance between these pseudo-bulk transcriptomes was driven by known parasite effector proteins **(Figure 2C and 2D)**. For example, the first principal component identifies a cluster of all known secreted *T. gondii* genes required for export of dense granule effectors (MYR1-4^13, 14, 36, 37^, ROP17^38^, GRA45^9, 37^, PPM3C^35^; GRA44 is also required but has a very strong negative growth phenotype so we did not recover enough cells to analyse^37, 39^), while the third principal component separates the dense granule effectors GRA16 and HCE1 from GRA24, indicating these effectors may drive opposing transcriptional responses in the host cell.

To formally identify these apparent clusters of functionally related effector proteins, we calculated Pearson correlation coefficients between the pseudo-bulk transcriptomes of the effectors that significantly perturb host cell gene expression **(Figure 2E)**. Hierarchical clustering based on these correlation coefficients identified four broad clusters of effector proteins. A cluster of tightly correlated pseudo-bulk transcriptomes contained six proteins that have been shown to be essential for dense granule effector export, and which likely constitute the “core” export machinery: MYR1-4, ROP17, and GRA45^9, 13, 36–38^. An additional cluster of dense granule proteins (comprising GRA59, GRA57, MAF1B, GRA25, PPM3C, GRA70) was moderately correlated with the cluster of essential export proteins, putatively indicating some similarity or overlap in phenotype. A third cluster of five genes contained ROP9, GRA7, and GRA3, together with the effectors HCE1 and GRA16 that have been shown to affect the host cell cycle and metabolism^15, 16, 19^. Finally, we identified a cluster of five genes we interpret as “immune regulators”: GRA28, which regulates host cell motility and cytokine expression through interaction with chromatin remodelers^21, 22^, GRA24, which activates p38α MAPK^20^, IST, which inhibits the induction of ISGs by recruiting repressive nucleosome remodellers to STAT1 target genes^17, 18^, ROP16, which induces M2 polarisation in host cells by directly phosphorylating and activating STAT3 and STAT6^23, 40, 41^, and an uncharacterised gene, TGGT1_222100.

To determine which host cell pathways are differentially regulated by these effector proteins, we scored the single cell transcriptomes for expression of 196 gene sets from the Pathway Interaction Database^42, 43^ using VISION^30^, and tested for up-or down-regulation of these pathways for each putative effector protein by Wilcoxon rank-sum test on the signature scores **(Figure 2F and S5, Supplementary Data 7)**. This analysis reaffirmed many of the known phenotypes of characterised effector proteins, for example upregulation of the c-Myc pathway by GRA16^44^, activation of p38ɑ MAP kinase by GRA24^20^, and suppression of interferon responses by IST^17, 18^. Furthermore, inspection of differentially regulated pathways suggested functions for the putative effector proteins newly identified here, such as activation of the E2F pathway by ROP9, indicating a role in regulation of the host cell cycle, and induction of an IL-4-like response by TGGT1_222100. Hierarchical clustering of the effectors using mean signature scores for 159 pathways significantly differentially regulated by at least one effector gave similar results to the clustering using Pearson correlation coefficients.

In summary, with dual perturb-seq we are able to screen >200 uncharacterised genes in intracellular *T. gondii* parasites for an effect on the transcriptome of mammalian host cells. With a single statistical test, we identify all currently known effector proteins that target host transcription, and identify six additional candidate effectors. By inspection of pseudo-bulk transcriptomes by PCA and inter-effector correlation, we identify clusters of functionally related effector proteins. Finally, we are able to determine which host cell pathways are differentially regulated by each effector protein, as a first step towards establishing their function.

### GRA59 contributes to export of dense granule effectors

One of the top hits in the dual perturb-seq screen was the dense granule protein GRA59 (TGGT1_313440) **(Figure 2B)**, for which no function is known^45^. Correlation of pseudo-bulk transcriptomes and clustering based on differentially expressed host cell pathways indicated that GRA59 has a similar host cell phenotype to genes that are known to be required for export of dense granule effectors into the host cell **(Figure 2E and 2F)**. By scoring sg(GRA59)-expressing cells for a ΔMYR1-like signature, we found that these cells had an intermediate phenotype compared to sg(MYR1)-expressing cells **(Figure 3A)**.

**Figure 3.**
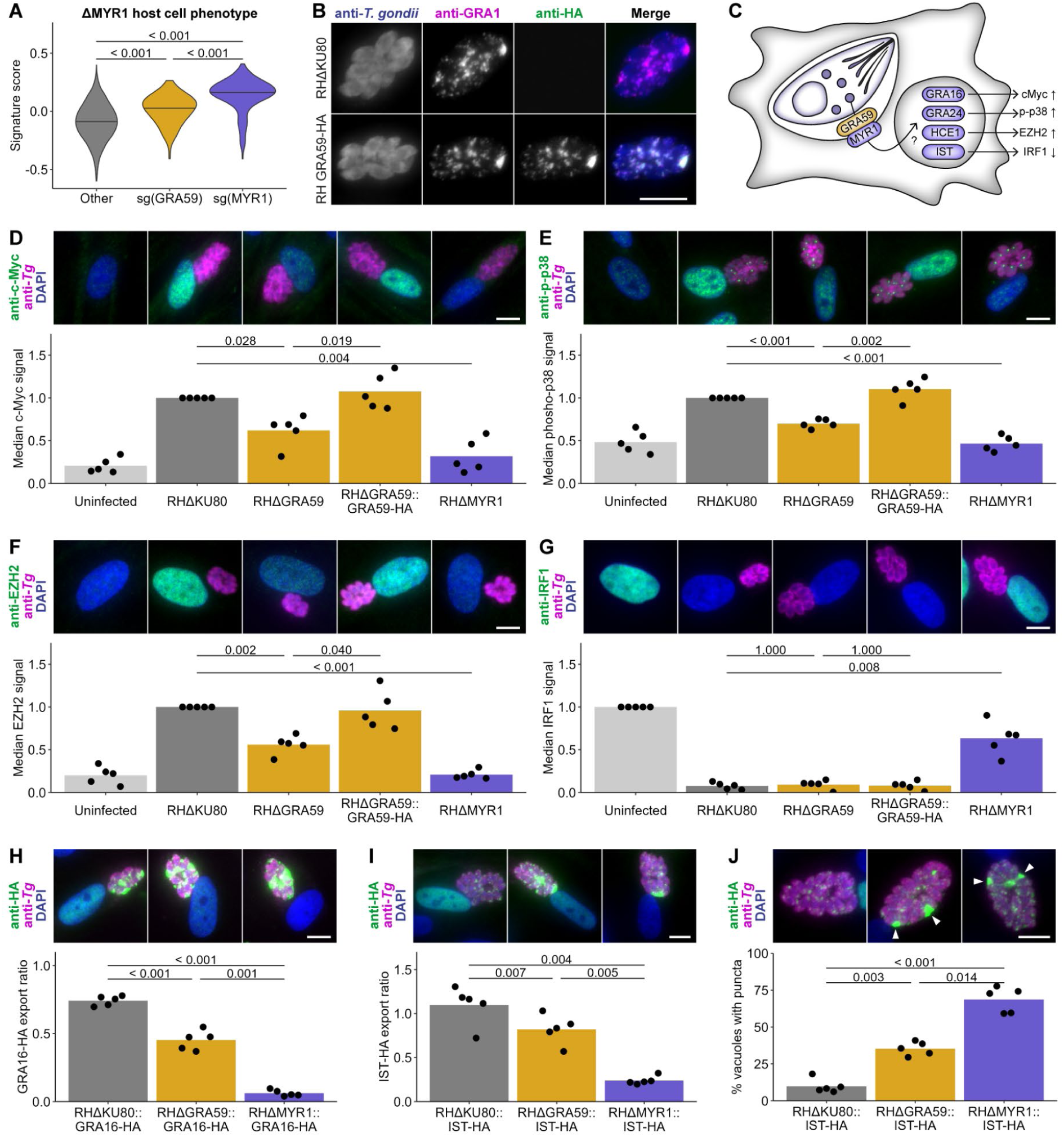
GRA59 (TGGT1_313440) contributes to dense granule effector protein export. **A.** Scoring of single cell transcriptomes for a ΔMYR1-infected phenotype using VISION. Differences tested by two-sided Wilcoxon rank sum test with Bonferroni adjustment. **B.** Immunofluorescence localisation of GRA59-HA. Scale bar = 10 µm. **C.** Model of dense granule effector export into the host cell. **D.** Quantification of nuclear c-Myc immunofluorescence in infected cells. Scale bar = 10 µm. Differences tested by two-sided Wilcoxon rank sum test with Bonferroni adjustment. **E.** Quantification of nuclear phospho-p38 immunofluorescence in infected cells. Scale bar = 10 µm. Differences tested by two-sided Wilcoxon rank sum test with Bonferroni adjustment. **F.** Quantification of nuclear EZH2 immunofluorescence in infected cells. Scale bar = 10 µm. Differences tested by two-sided Wilcoxon rank sum test with Bonferroni adjustment. **G.** Quantification of nuclear IRF1 immunofluorescence in infected cells stimulated with IFNγ. Scale bar = 10 µm. Differences tested by two-sided Wilcoxon rank sum test with Bonferroni adjustment. **H.** Ratio of GRA16-HA immunofluorescence in the host cell nuclei compared to the vacuole. Scale bar = 10 µm. Differences tested by paired two-sided *t*-test with Bonferroni adjustment. **I.** Ratio of IST-HA immunofluorescence in the host cell nuclei compared to the vacuole. Scale bar = 10 µm. Differences tested by paired two-sided *t*-test with Bonferroni adjustment. **J.** Percentage of vacuoles containing accumulations of IST-HA immunofluorescence. White arrowheads indicate accumulations. Scale bar = 10 µm. Differences tested by paired two-sided *t*-test with Bonferroni adjustment.

We tagged GRA59 with a single HA epitope at the C-terminus **(Figure S6A and S6B)** and confirmed prior immunofluorescence localisation to the parasitophorous vacuole^45^ **(Figure S6C-E)**. In fully permeabilised cells, GRA59 co-localised with the vacuole lumen protein GRA1 and the intravacuolar network (IVN)-resident GRA2 **(Figure 3B, S6C, and S6D)**. However, upon partial permeabilisation with saponin, which disrupts the host cell plasma membrane but leaves the parasitophorous vacuole membrane (PVM) mostly intact, anti-HA staining was detectable that partially co-localised with the PVM marker GRA3, indicating that the C-terminus of GRA59 may be exposed to the host cell cytosol **(Figure S6E)**. In support of this finding, GRA59 was recently shown to be accessible to a host cell-localised biotin ligase^46^. We therefore hypothesised that GRA59 may be a component of the export complex in the PVM **(Figure 3C)**.

We generated GRA59 knockout and complemented cell lines in the RHΔKU80 background **(Figure S7)** and measured dense granule effector export by quantifying nuclear c-Myc immunofluorescence of infected HFFs. Upregulation of host c-Myc by *T. gondii* is entirely dependent on MYR1-dependent export^13^ and partially dependent on the exported effector GRA16^44^. While the parental RHΔKU80 strain strongly induced host c-Myc and knockout of MYR1 completely ablated this upregulation, as expected, knockout of GRA59 resulted in an intermediate phenotype that was restored by complementation **(Figure 3D)**. This phenotype is consistent either with a role for GRA59 in effector export, or in directly upregulating c-Myc in addition to or in partnership with GRA16. We therefore measured three additional host cell phenotypes of MYR-dependent exported effector proteins: phosphorylation and nuclear translocation of p38 MAP kinase by GRA24^20^, upregulation of EZH2 by HCE1/TEEGR^15, 16^, and inhibition of IFNγ-induced IRF1 expression by IST^17, 18^. For both p38 phosphorylation and EZH2 upregulation, knockout of GRA59 resulted in a phenotype intermediate between the parental RHΔKU80 strain and RHΔMYR1 similarly to c-Myc upregulation **(Figure 3D and 3E)**. These results indicate that GRA59 contributes to the export of multiple effector proteins, but that knockout does not completely abolish export, as is the case for MYR1. Surprisingly, suppression of IFNγ-mediated IRF1 induction was not affected by GRA59, suggesting normal export of the effector protein IST **(Figure 3G)**.

To directly test whether effector export was affected by GRA59, we introduced an HA-tagged variant of either GRA16 or IST to the RHΔKU80, RHΔGRA59, and RHΔMYR1 strains **(Figure S8)**. Following export to the host cell, both GRA16 and IST localise to the host cell nucleus. Therefore, we quantified the export of GRA16-HA and IST-HA as the ratio of anti-HA immunofluorescence signal in the host cell nuclei compared to that remaining in the vacuole at 24 hpi. In accordance with the c-Myc phenotype, we observed an intermediate reduction in GRA16-HA export by the RHΔGRA59 strain compared to RHΔKU80 and RHΔMYR1 **(Figure 3H)**. Surprisingly, export of IST-HA was also reduced in the RHΔGRA59 strain, even though suppression of IRF1 was not affected **(Figure 3I)**. Furthermore, we noticed the presence of accumulations of IST-HA signal within the vacuole of the RHΔGRA59 and RHΔMYR1 strains that were largely absent in RHΔKU80 **(Figure 3J)**. The percentage of vacuole containing these IST-HA accumulations mirrored the reduced export to the host cell nucleus, indicating retention of IST-HA within the vacuole when export is limited. Nearly all vacuoles, including those of RHΔKU80, contained accumulations of GRA16-HA, therefore it was not possible to quantify retention of GRA16-HA in the same manner.

Together, these data confirm the apparent phenotype of GRA59 in the dual perturb-seq screen: GRA59 has a MYR1-like phenotype and contributes to the export of dense granule proteins from the vacuole into the host cell. However, unlike for MYR1, knockout of GRA59 does not completely ablate export. Instead, there is an intermediate reduction in export of GRA16 and IST, and indications that export of GRA24 and HCE1 are similarly affected.

### SOS1 is required for sustained STAT6 signalling

Aside from established effectors, the gene with the highest perturbation score in the screen was the uncharacterised gene TGGT1_222100 **(Figure 2B)**, which, based on the following results, we propose naming *Tg*SOS1. The pseudo-bulk transcriptome of sg(SOS1)-expressing cells correlated most strongly with the rhoptry effector kinase ROP16 **(Figure 2E)**, which has been shown to directly phosphorylate STAT3 and STAT6 to activate downstream transcriptional programmes^23, 41^. Correspondingly, the most strongly downregulated genes in sg(SOS1)-expressing cells were CCL11 and CCL26, both of which have been shown to be regulated by STAT6^47, 48^ and were previously identified among the most strongly downregulated genes in *T. gondii* RHΔROP16-infected HFFs compared to RH wild type-infected HFFs (Ong, Reese, and Boothroyd 2010) **(Figure 4A)**. Similarly, the most strongly downregulated pathway in these cells was the response to IL-4, which is the canonical activator of STAT6 signalling^49–52^ **(Figure 4B, Supplementary Data 7)**.

**Figure 4.**
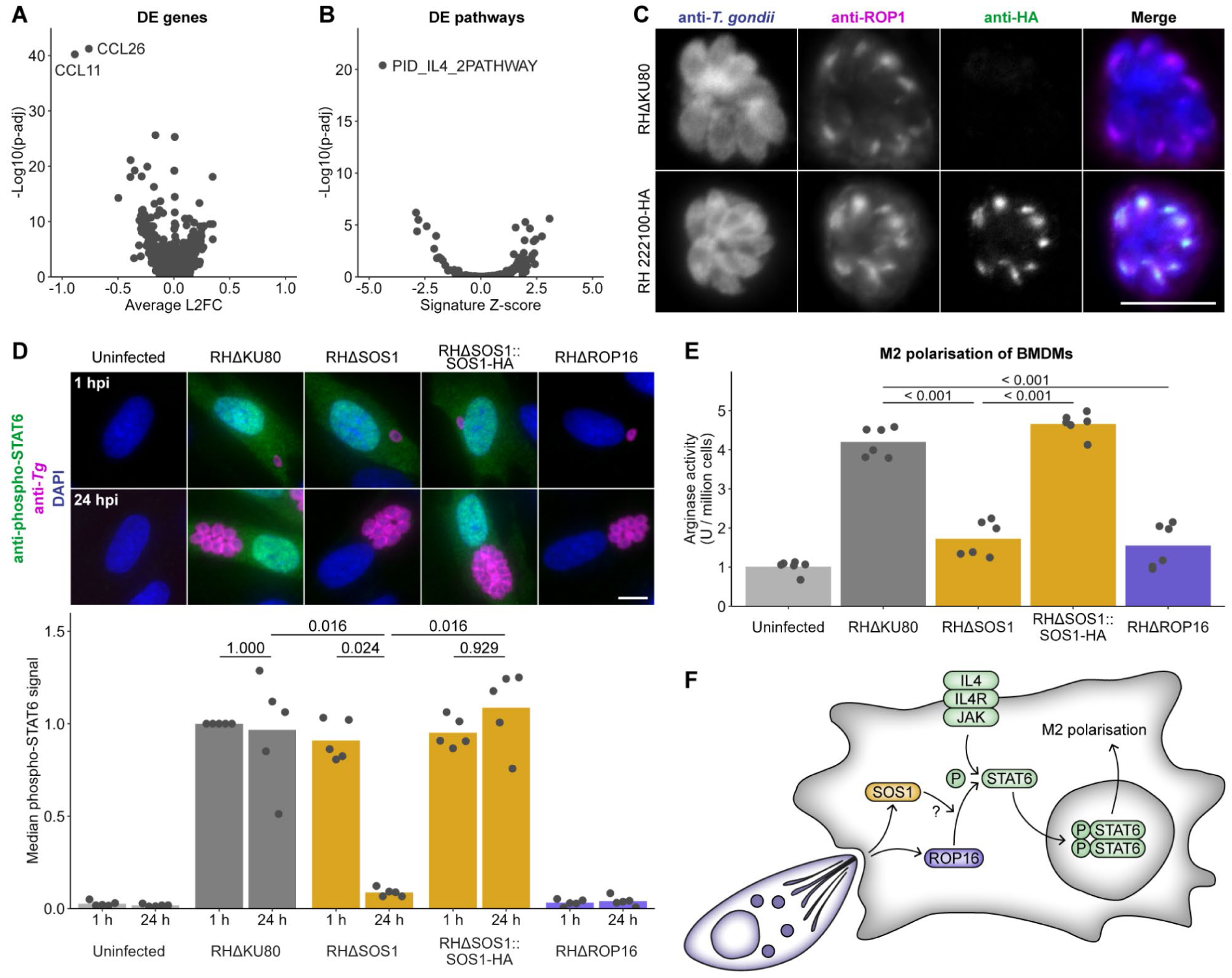
SOS1 (TGGT1_222100) is required for sustained STAT6 signalling in infected cells. **A.** Differentially expressed host cell genes for sg(SOS1)-expression cells compared to all other (non-control) cells (two-sided Wilcoxon rank-sum test with Benjamini-Hochberg adjustment). **B.** Differentially expressed Pathway Interaction Database gene sets for sg(SOS1)-expression cells compared to all other (non-control) cells (two-sided Wilcoxon rank-sum test with Benjamini-Hochberg adjustment). **C.** Immunofluorescence localisation of SOS1-HA. Scale bar = 10 µm. **D.** Quantification of nuclear phospho-STAT6 immunofluorescence in infected HFFs at 1 h and 24 h post-infection. Scale bar = 10 µm. Differences tested by two-sided Wilcoxon rank sum test with Bonferroni adjustment. **E.** Arginase activity of infected cell lysate. Differences tested by two-sided *t*-test with Bonferroni adjustment. **F.** Model of ROP16 and SOS1-dependent STAT6 signalling.

SOS1 co-localises with ROP1 at the rhoptry bulb **(Figure 4C)**. SOS1 has a predicted molecular weight of 128 kDa and appears to be cleaved approximately in the middle, with the tagged C-terminal fragment migrating at 70 kDa **(Figure S9B)**. This processing could be inhibited with the small molecule 49c, an inhibitor of the Golgi-resident aspartyl protease ASP3^53^ **(Figure S9D)**. This result indicates that, despite the apparent absence of a signal peptide, SOS1 is within the secretory pathway and contained within the lumen of the rhoptry bulb. This putative ASP3 cleavage site lies within a predicted flexible linker between two uncharacterised domains **(Figure S9E)**. Unusually for a rhoptry protein, the predicted C-terminal domain is conserved across Apicomplexa, while the N-terminal domain is recognisable only in Coccidia.

Based on the apparent similarity of sg(SOS1)-expressing cells to sg(ROP16)-expressing cells, we hypothesised that knockout of SOS1 would phenocopy knockout of ROP16 in loss of host STAT6 phosphorylation. We generated RHΔSOS1 and RHΔROP16 knockout cell lines, complemented RHΔSOS1 by inserting an HA-tagged copy of SOS1 **(Figure S10)**, and measured phospho-STAT6 immunofluorescence in HFFs infected with these strains. Surprisingly, we did not find any difference in STAT6 phosphorylation at 1 hpi for RHΔSOS1 compared to RHΔKU80, despite observing the expected loss of this phosphorylation in cells infected with the RHΔROP16 cell line **(Figure 4D)**. However, at 24 hpi, the same time point at which the dual perturb-seq data were collected, we found that STAT6 signalling was completely abolished in the RHΔSOS1-infected cells and rescued by complementation. Therefore, while ROP16 is sufficient for the initial phosphorylation of STAT6, SOS1 is required to sustain STAT6 signalling throughout the infection. Hence, we name this gene (TGGT1_222100) **S**ustainer **O**f **S**TAT signalling **1**.

Activation of STAT6 signalling by ROP16 induces M2 polarisation of *T. gondii*-infected macrophages, which is implicated in dampening anti-parasitic immune responses^54, 55^. To test if SOS1-dependent sustained STAT6 phosphorylation is important for M2 polarisation, we infected murine bone marrow-derived macrophages (BMDMs) and measured arginase activity, a defining feature of M2 polarisation^56^. Arginase activity was strongly induced by infection with RHΔKU80, but was reduced to near the level in uninfected cells by knockout of either ROP16 or SOS1 **(Figure 4E)**.

These results show that the rhoptry kinase ROP16 is not sufficient for sustained host cell reprogramming, and reveal a requirement for the previously unknown effector protein, SOS1. While ROP16 is required for the initial phosphorylation of STAT6, SOS1 is necessary to sustain this signalling and thereby facilitate durable M2 polarisation of infected macrophages **(Figure 4F)**.

### SOS1 maintains efficient bradyzoite cyst formation in neurons

During acute infection *in vivo*, a small percentage of *T. gondii* tachyzoites differentiate into bradyzoites, forming tissue cysts that are highly resistant to antiparasitic drugs and to clearance by the host immune system. Bradyzoite cysts can persist for the lifetime of the host and are essential in the *T. gondii* life cycle for transmission to the definitive felid host^57^; moreover, they are clinically important as a source of recrudescent infection in immunocompromised individuals^58^. ROP16-dependent STAT6 signalling has previously been shown to facilitate bradyzoite cyst formation *in vivo* and *in vitro*, although the downstream host cell pathways involved remain unknown^59, 60^. The findings here for SOS1 raised the question of whether sustained STAT6 signalling is necessary to maintain a permissive host cell environment for cyst formation, or whether only the initial ROP16-dependent burst of signalling is necessary to trigger cyst formation.

As the *T. gondii* RH strain used in this work so far exhibits a very low rate of differentiation to bradyzoites^61^, to investigate the role of SOS1 in cyst formation we generated an SOS1 knockout in the type III *T. gondii* CEP strain **(Figure S11)**. As in the RH strain, we found that infection of HFFs with CEPΔSOS1 induced an initial ROP16-dependent burst of STAT6 signalling that dissipated after 24 h **(Figure 5A)**. As *T. gondii* forms bradyzoite cysts primarily in neurons and skeletal muscle *in vivo*, we also measured STAT6 phosphorylation in infected primary murine neurons. We observed the same loss of STAT6 phosphorylation at 24 hpi in neurons infected with CEPΔSOS1, compared to the parental strain, as we observed in HFFs, confirming that this phenotype is independent of host cell type **(Figure 5B)**.

**Figure 5.**
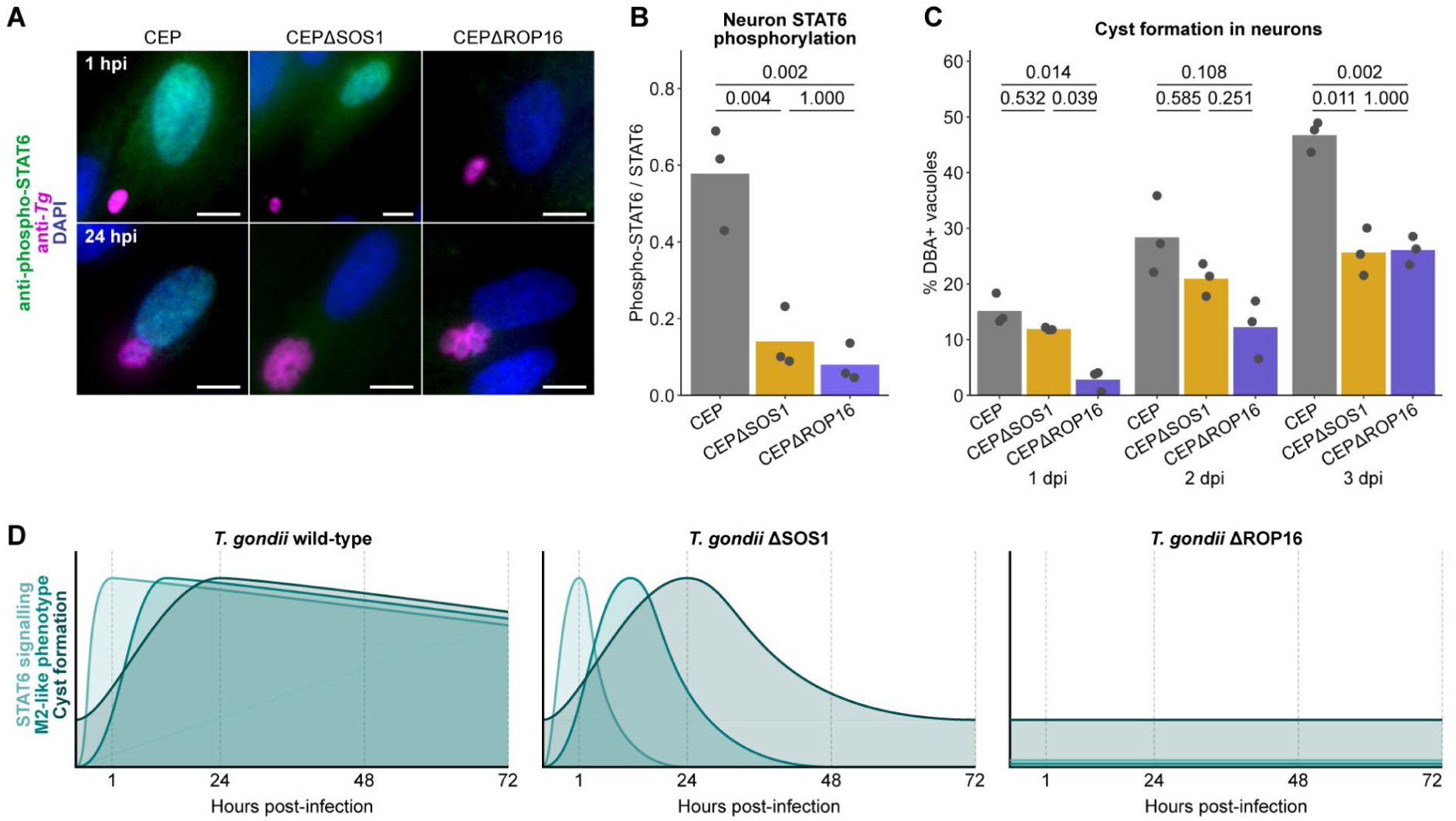
SOS1 maintains efficient bradyzoite cyst formation in neurons. **A.** Phospho-STAT6 immunofluorescence in infected HFFs at 1 h and 24 h post-infection. Scale bar = 10 µm. **B.** Quantification of phospho-STAT6 relative to total STAT6 from Western blots of samples of primary murine neurons infected for 24 h with indicated *T. gondii* strains. Differences tested by two-sided *t-*test with Bonferroni adjustment. **C.** Percentage of DBA-positive bradyzoite cysts out of total vacuoles with ≥2 parasites in primary murine neurons infected for 1, 2 or 3 days. Differences test by two-sided *t-*test with Bonferroni adjustment. **D.** Model of ROP16 and SOS1-mediated host cell reprogramming.

To determine whether SOS1 is required for efficient cyst formation, as is ROP16, we used a previously described high-content imaging and automated image analysis to quantify parasite differentiation from tachyzoites to bradyzoites in primary murine neurons^60^. Conversion of tachyzoite vacuoles to bradyzoite cysts was determined by staining of the cyst wall with *Dolichos bisflorus* agglutinin (DBA) (**Figure 5C)**. At 24 hpi, there was no difference in the percentage of DBA-positive vacuoles between the parental CEP strain and CEPΔSOS1. However, at later timepoints, the rate of bradyzoite conversion in the CEPΔSOS1 line appeared to decrease, until at 72 hpi the percentage of DBA positive vacuoles was significantly reduced compared to the parental CEP stain and indistinguishable from CEPΔROP16. This finding mirrors the dependence of sustained STAT6 signalling and M2 polarisation on SOS1, showing that SOS1 is also critical for efficient cyst formation **(Figure 5D)**.

## DISCUSSION

In this work, we combined pooled knockout screening with multi-modal single-cell RNA-seq to overcome prior challenges in identifying *T. gondii* effector proteins that manipulate host cell gene expression. We propose this dual perturb-seq as a method to investigate transcriptional interactions in any host-pathogen or host-symbiont models that are amenable to genetic manipulation using CRISPR-Cas systems, with only minor modifications to the experimental strategy and analysis described here being necessary. For example, here we rely on non-homologous end-joining to induce loss-of-function mutations, a pathway which is absent in many protozoan parasites. However, this requirement could be obviated by replacement of Cas9 in the perturb-seq vector with catalytically inactive Cas9, for CRISPRi, or Cas13, to target RNA transcripts. Furthermore, whereas sgRNAs are typically delivered by lentiviral transduction, we instead deliver both Cas9 and the sgRNA simultaneously by electroporation with a plasmid vector, highlighting the feasibility of perturb-seq in diverse non-mammalian cells.

For *T. gondii*, a ubiquitous and clinically significant pathogen, we were able to use dual perturb-seq data to infer the functions of previously uncharacterised genes *de novo* and in relation to established effector proteins. Importantly, despite critical roles in infection *in vivo*, effector proteins that target host cell transcription have not been identified by growth competition-based knockout screens *in vitro* and *in vivo*^7–10^. This is likely because the primary role of these effectors in modulating host immunity is rescued in a pooled infection^7^. This phenomenon necessitated the use of dual perturb-seq to directly measure the impact of *T. gondii* gene knockout on the host cell transcriptome in a high-throughput and unbiased manner.

We validated the dual perturb-seq phenotype of GRA59 by showing that it contributes to export of other dense granule effectors into the host cell. Based on these phenotypes, plus the localisation and topology of GRA59, we suggest that it may be an accessory or regulatory component of the export complex, facilitating efficient effector export into the host cell. Interestingly, we do not identify any mutants that completely ablate effector export other than the “core” cluster comprising MYR1-4, ROP17, and GRA45. Having thus defined all components required for protein export paves the way for detailed mechanistic and structural studies of this pathway, which is currently poorly understood yet critical for *T. gondii* virulence^5, 13^.

Additionally, we identify a previously unknown effector protein, SOS1, which operates via a novel mode of action to prolong STAT6 signalling initiated by the ROP16 kinase. Such a finding was unexpected, as recombinant ROP16 is sufficient to phosphorylate STAT6 *in vitro*^40, 41^. In a cellular context, SOS1 may regulate the activity or stability of ROP16, or may disrupt feedback mechanisms that downregulate STAT6 activity. We show here that sustained STAT6 signalling is necessary for M2 polarisation of *T. gondii* infected macrophages and for efficient bradyzoite cyst formation in neurons. Through these phenotypes, SOS1 is implicated in parasite immune evasion and transmission *in vivo*.

With dual perturb-seq, we are thus able to identify effector proteins in a high-throughput and systematic manner and reveal new mechanisms of effector-mediated host cell reprogramming. Dual perturb-seq will be a powerful tool to accelerate the study of *T. gondii* effector proteins across different strains and host cell types, and to identify the molecular mediators of transcriptional remodelling in other host-microbe interactions.

## METHODS

### Mice

Wild-type C57BL/6J mice were bred and housed in pathogen-free conditions at the Biological Research Facility of the Francis Crick Institute in accordance with the Home Office UK Animals (Scientific Procedures) Act 1986 and European Union Directive 2010/63/EU. No regulated procedures were carried out on live animals at the Francis Crick Institute in this work.

Ai6 Cre reporter mice^62^ were bred and housed in specific-pathogen-free conditions at the University of Arizona Animal Care facilities with a 14 hr/10 hr light/dark cycle, with ambient temperature between 68° and 75° F, and 30–70% humidity. All procedures and experiments were carried out in accordance with the Public Health Service Policy on Human Care and Use of Laboratory Animals and approved by the University of Arizona’s Institutional Animal Care and Use Committee (#12-391).

### Cell culture

#### HFF

Human foreskin fibroblasts (ATCC, SCRC-1041) were cultured in Dulbecco’s Modified Eagle’s Medium with 4.5 g/L glucose and

GlutaMAX (Gibco) supplemented with 10% heat-inactivated foetal bovine serum (Gibco) at 37 °C and 5% CO2.

#### BMDM

Monocytes isolated from the bone marrow of 6-12-week-old male C57BL6/J mice were differentiated into macrophages for six days in 70% RPMI 1640 ATCC modification (Gibco), 20% L929 cell-conditioned medium (provided by the Cell Services Science Technology Platform at the Francis Crick Institute), 10% heat-inactivated FBS (Gibco), 100 U/mL penicillin-streptomycin (Gibco) and 50 μM 2-mercaptoethanol (Sigma). Following differentiation, 2-mercaptoethanol was removed from the medium.

#### Primary neurons

Primary murine neuronal cell cultures were derived from Ai6 Cre reporter mice as previously described^63^. Briefly, cortical neurons were dissected from E17 embryos, dissociated, and seeded in poly-L-lysine-coated 96-well, clear-bottom, black-walled plates at a density of 20,000 cells per well in Minimal Essential Medium (Thermo) supplemented with D-glucose, L-glutamine, and 5% FBS. After four hours, the medium was changed to Neurobasal Medium (Thermo) with B-27 Supplement (Thermo), L-glutamine, and penicillin-streptomycin. After four days of *in vitro* culture, neurons received a half-volume medium change with complete Neurobasal Medium additionally supplemented with 5 µM cytosine arabinoside to prevent glial proliferation. Thereafter, one third medium exchanges with complete Neurobasal Medium occurred every 3-4 days. Experiments were carried out using the neurons following 10-12 days of *in vitro* culture.

#### Toxoplasma gondii

*T. gondii* tachyzoites were maintained by serial passage in HFFs every 2-3 days and isolated for experiments by passing through a 27-gauge needle followed by a 5 µm filter. Parental strains used in this study were RHΔHXGPRT^64^, RHΔKU80^65^, and CEPΔHXGPRT^59^. Parasite genotype was verified by restriction fragment length polymorphism of the SAG3 gene as previously described^66^. All *T. gondii* strains were regularly checked for *Mycoplasma spp.* contamination by PCR.

### Generation of perturb-seq vectors

Capture sequences 1 and 2 were inserted into the pCas9-sgRNA vector^7^ by inverse PCR using primers 1-4 **(Supplementary Data 8)**. To generate individual perturb-seq vectors, the protospacer was modified by inverse PCR using primers 5-29 **(Supplementary Data 8)**. To generate the pool of perturb-seq vectors used in the screen, ssDNA oligonucleotides encoding the protospacer sequences were selected from an arrayed library^7^ using an Echo 550 Acoustic Liquid Handler (Labcyte). Oligonucleotides were dispensed in triplicate on three different days to reduce loss due to incomplete thawing of frozen stocks or misalignment of the plates. The pooled oligonucleotides were integrated into the pCas9-sgRNA-CS1 vector by Gibson assembly as previously described^7^. An Illumina sequencing library was prepared from the resulting plasmid pool to verify incorporation of the protospacer sequences by nested PCR using primers 30-33 **(Supplementary Data 8)**. The resulting library was sequenced on a HiSeq 4000 platform (Illumina) with 100 bp paired-end reads to a depth of 30 million reads **(Supplementary Data 3)**. The numbers of reads matching each protospacer sequence were counted using a custom perl script.

### Dual perturb-seq experiments

*T. gondii* RHΔHXGPRT were transfected with KpnI-HF-digested (NEB) perturb-seq vectors by electroporation using the Amaxa 4D Nucleofector system (Lonza) with buffer P3 and pulse code EO-115. For individual perturb-seq vectors, at least 10^7^ parasites were transfected with 15 µg of plasmid in one well of a 4D Nucleofector X Unit 16-well strip. For the perturb-seq plasmid pool, at least 10^8^ parasites were transfected with 120 µg of plasmid split across eight wells of a 16-well strip. 24 h after transfection, 25 μg/mL mycophenolic acid (Sigma) and 50 μg/mL xanthine (Sigma) were added to the parasites to select for integration of the linearised perturb-seq vector into the genome for at least seven days.

24 h prior to infection, the HFF medium was changed to Dulbecco’s Modified Eagle’s Medium with 4.5 g/L glucose and GlutaMAX supplemented with 2% heat-inactivated FBS. The HFFs were infected with transfected parasites at a MOI of 0.1. After 1 h the medium was changed to remove remaining extracellular parasites. Where IFNγ stimulated was used, recombinant human IFNγ (BioTechne) was added to the infected cells at 21 h post-infection to a final concentration of 5 ng/mL (∼10 U/mL).

24 h post-infection, the HFFs were dissociated using TrypLE Express (Gibco) and stained with 5 µg/mL DAPI (Sigma) for 5 minutes on ice. The cells were pelleted by centrifugation at 300 x g for 5 minutes at 4 °C, resuspended in 2% FBS DMEM without phenol red and passed through a 30 µm filter. Infected cells were enriched based on Cas9-GFP expression using a FACSAria III cell sorter (BD).

The sorted cells were concentrated by centrifugation at 300 x g for 5 minutes at 4 °C and resuspended in DPBS (Sigma) with 0.04% w/v BSA (Sigma). The cells were partitioned into gel bead-in-emulsion droplets for barcoding and reverse transcription using the Chromium Controller (10x Genomics) and Single Cell 3’ GEM, Library & Gel Bead Kit v3.1 (10x Genomics) according to the manufacturer’s instructions, with a targeted recovery of 10,000 cells per channel. 3’ mRNA and CRISPR sgRNA sequencing libraries were constructed following the manufacturer’s protocol for 3’ gene expression with Feature Barcoding technology for CRISPR screening, using the additional Chromium Single Cell 3ʹ Feature Barcode Library Kit. The libraries were sequenced on a HiSeq 4000 platform (Illumina) with a targeted depth of 50,000 reads per cell for the 3’ mRNA library and 5000 reads per cell for the CRISPR sgRNA library.

### Analysis of dual perturb-seq data

A schematic overview of the analysis can be found in **Figure S1**.

#### Pre-processing, filtering, and normalisation

A multiple species reference genome comprising *Homo sapiens* GRCh38 release 95 and *Toxoplasma gondii* ME49 TGA4 release 42 was generated using Cell Ranger v3.0.2 (10x Genomics). Using Cell Ranger v6.1.2, sequencing reads from the 3’ mRNA and CRISPR sgRNA libraries were aligned to this reference and a custom reference of protospacer sequences, including intronic reads but leaving all other settings as the default. All subsequent analyses were carried out in R v4.0.2 (www.r-project.org). UMI count matrices from Cell Ranger were imported into Seurat v4.2.0^67^ with *H. sapiens* mRNA, *T. gondii* mRNA, and CRISPR sgRNA counts split into separate assays of a single Seurat object. Cell barcodes were filtered to retain only cells with a high total count of both *H. sapiens* and *T. gondii* mRNA UMIs, corresponding to infected cells. Cells with >10% UMIs derived from mitochondrial genome-encoded transcripts, characteristic of low-quality cells^68^, were removed. Finally, we filtered the cells to retain only those in which a single protospacer sequence was detected, with the aim of eliminating multiplet droplets or HFFs infected by multiple parasites. The remaining cells were classified based on the *T. gondii* gene target of the detected protospacer sequence. *H. sapiens* mRNA counts were normalised using SCTransform v2^69, 70^. Replicates (Chromium channels) of the perturb-seq screen from both unstimulated and IFNγ-stimulated samples were merged and normalised using SCTransform v2, passing the replicate ID as a variable to regress out of the SCTransform residuals.

#### Identification of effector proteins

The SCTransform residuals (“scale.data” slot) of the top 3000 most variable genes were analysed by principal component analysis using Seurat. For each *T. gondii* target gene, the distribution of infected cells in the top 20 principal components was compared to all other cells (excluding cells infected with known effector proteins) using Hotelling’s *t*^2^-test as implemented in the Hotelling v1.0.8 package. Hits were called based on a Benjamini-Hochberg-adjusted *p-*value of 0.01 or less.

#### Pseudo-bulk analyses

For each *T. gondii* target gene represented by at least 30 single cells, pseudo-bulk transcriptomes were generated from the mean SCTransform residuals of the top 3000 most variable genes. These pseudo-bulk profiles were analysed by principal component analysis. For effectors identified by Hotelling’s *t*^2^-test, Pearson correlation coefficients were calculated between the pseudo-bulk transcriptomes and clustered using the complete linkage method implemented in the hclust function of the stats package v4.0.4.

#### Identification of differentially expressed genes

In the pilot experiments, differentially expressed host genes were identified by two-sided Wilcoxon rank-sum test on the SCTransform-normalised data using the sg(UPRT)-expressing cells as a reference population. In the screen, differentially expressed host genes were identified for each effector using all other cells (excluding controls) as a reference population. *p*-values were adjusted using the Benjamini-Hochberg procedure.

#### Identification of differentially expressed pathways

Single cells were scored for the expression of gene sets in Pathway Interaction Database^42, 43^ using the SCTransform-normalised gene expression data with VISION v2.1.0^30^. VISION signature scores were converted to Z-scores and, for each *T. gondii* target gene represented by at least 30 single cells, up-or down-regulation of each pathway in cells infected with that knockout compared to all other knockouts (excluding known effector proteins) was tested by two-sided Wilcoxon rank-sum test. For the effector proteins identified by Hotelling’s *t*^2^-test, the mean pathway Z-score was plotted for each of the pathways differentially expressed by at least one effector (Benjamini-Hochberg-adjusted *p*-value ≤0.05). The effectors and gene sets were clustered using these mean Z-scores using the complete linkage method.

#### Comparison to published bulk phenotypes using VISION signature scores

Host cell genes differentially expressed upon infection with a *T. gondii* knockout strain compared to a wild-type strain were extracted from bulk RNA-seq/microarray data for seven effector proteins: MYR1 (6 hpi RHΔMYR1-infected HFFs)^14^, HCE1 (6 hpi RHΔHCE1-infected HFFs)^15^, IST (6 hpi RHΔIST-infected HFFs)^71^, GRA16 (18-20 hpi RHΔGRA16-infected HFFs)^19^, GRA24 (18-20 hpi RHΔGRA24-infected murine BMDMs)^20^, GRA28 (RHΔGRA28-infected THP-1s, timepoint not stated)^21^, and ROP16 (24 hpi type II::ROP16^type^ ^I^ versus type II wild-type-infected HFFs)^23^. Single cell transcriptomes were scored for the expression of these gene sets using the SCTransform-normalised gene expression data with VISION v2.1.0^30^, resulting in a “signature score” for each marker gene set for each cell. A higher signature score indicates that the same genes are up-or down-regulated in the single cell transcriptome data as in bulk transcriptome data. *T. gondii* knockout-infected cells were tested for higher scores of the corresponding signature compared to UPRT knockout (pilot experiments) or all other cells (screen experiment) by Wilcoxon rank-sum test.

### Generation of *T. gondii* cell lines

#### C-terminal epitope tagging

Cas9-sgRNA plasmids (without HXGPRT)^72^ targeting the 3’ UTRs of GRA59 and SOS1 were generated by inverse PCR using primers 5, 34, and 35 **(Supplementary Data 8)**. In-frame HA-Ter^GRA2^::Pro^DHFR-^HXGPRT-Ter^DHFR^ repair constructs were amplified from a plasmid template ^10^ using primers 36-39 to introduce 40 bp homology arms to the 3’ UTRs of GRA59 and SOS1. For each strain, 15 µg each of Cas9-sgRNA plasmid and repair construct were co-transfected into the *T. gondii* RHΔKU80 by electroporation as described above. 24 h after transfection, 25 μg/mL mycophenolic acid (Sigma) and 50 μg/mL xanthine (Sigma) was added to the culture medium for at least seven days to select for integration of the repair construct into the genome, following which clonal cell lines were obtained by limiting dilution. Integration of the repair construct into the genome was verified by diagnostic PCR using primers 40-43 on parasite genomic DNA extracted using the DNeasy Blood and Tissue Kit (Qiagen).

#### Knockouts (T. gondii RH strain)

Cas9-sgRNA plasmids (without HXGPRT) targeting the coding sequence of the target gene were generated by inverse PCR using primers 5 and 44-47 **(Supplementary Data 8)**. Pro^GRA1^-mCherry-T2A-HXGPRT-Ter^GRA2^ repair constructs were amplified from a template plasmid using primers 48-55 to introduce 40 bp homology arms to the 5’ and 3’ UTRs of the target gene. For each target gene, 15 µg each of Cas9-sgRNA plasmid and repair construct were co-transfected into *T. gondii* RHΔKU80 by electroporation as described above. 24 h after transfection, 25 μg/mL mycophenolic acid (Sigma) and 50 μg/mL xanthine (Sigma) was added to the culture medium for at least seven days to select for integration of the repair construct into the genome, following which clonal cell lines were obtained by limiting dilution. Integration of the repair construct into the genome was verified by diagnostic PCR using primers 56-63.

#### Knockouts (T. gondii CEP strain)

A Cas9-sgRNA (without HXGPRT) with two sgRNAs targeting the 5’ and 3’ UTRs of SOS1 was generated as previously described^73^ using primers 77-80 to introduce the protospacer sequences by inverse PCR **(Supplementary Data 8)**. A repair plasmid to replace the coding sequence of SOS1 with an HXGPRT-mCherry construct was generated by amplifying the 5’ and 3’ UTRs of SOS1 (adjacent to the protospacers in the Cas9-sgRNA plasmid) using primers 81-84 and inserting the resulting fragments into the pTKO2 plasmid^27^ by In-Fusion cloning (Takara). Parasites were co-transfected with both the Cas9-sgRNA and pTKO2 plasmids, selected with mycophenolic acid and xanthine, as clonal cell lines obtained as described above. Deletion of the SOS1 CDS was verified by diagnostic PCR on extracted genomic DNA using primers 85-86, with amplification of the SAG1 CDS with primers 87-88 used as a control. Loss SOS1 mRNA expression relative to actin mRNA was verified by quantitative real-time PCR using primers 89-92.

#### Complementation

The coding sequence and 1000-1500 bp upstream of the start codon of the target gene were amplified from RHΔKU80 genomic DNA using primers 64-71 **(Supplementary Data 8)** and integrated into the pUPRT plasmid^72^ by Gibson assembly. 15 µg of pUPRT plasmid was linearised with AgeI-HF (NEB) (GRA59, SOS1, IST) or ScaI-HF (NEB) (GRA16) and co-transfected with 15 µg Cas9-sgRNA plasmid (without HXGPRT) targeting the UPRT locus. 24 h after transfection, 5 μM 5-fluorodeoxyuridine was added to the culture medium for at least seven days to select for integration of the pUPRT vector into the UPRT locus, following which clonal cell lines were obtained by limiting dilution. Integration of the pUPRT vector into the UPRT locus was verified by diagnostic PCR using primers 72-76.

### Western blot

#### HA-tagged parasite cell lines

Parasites were isolated by syringe-lysis, filtering, and washing in DPBS then lysed in RIPA buffer (Thermo) supplemented with cOmplete Mini EDTA-free Protease Inhibitor Cocktail (Roche) and 2 U/mL benzonase nuclease (Sigma) for 30 min on ice. Protein concentration was quantified using the Pierce BCA protein assay kit (Thermo). The samples were centrifuged at 15,000 x g for 5 min at 4 °C and the pellet discarded. 10 μg protein per sample was heated to 95 °C for 5 min in sample loading buffer and separated by SDS-PAGE using the Mini-PROTEAN electrophoresis system (Bio-Rad). Proteins were transferred to a nitrocellulose membrane using the Trans-Blot Turbo transfer system (Bio-Rad), blocked in 2% w/v skim milk powder, 0.1% v/v Tween 20 in PBS for 1 h at room temperature. The membrane was incubated in 1:1000 rat anti-HA (Sigma) or 1:200 mouse anti-*T. gondii* (Santa Cruz) for 1 h at room temperature followed by 1:10,000 goat anti-rat IRDye 800CW (LI-COR) or 1:10,000 goat anti-mouse IRDye 680LT (LI-COR) for 1 h at room temperature. Images were acquired using an Odyssey infra-red imaging system (LI-COR).

#### STAT6 phosphorylation in primary neurons

Primary neurons were infected for 24 h then protein was extracted by sonication in RIPA buffer supplemented with Protease Inhibitor Cocktail (Sigma) and Phosphatase Inhibitor Cocktail 2 (Sigma), as previously described^74^. 30 μg protein per sample was separated by SDS-PAGE, transferred to a PVDF membrane, and blocked as above. The membrane was stained with rabbit-anti STAT6 (Cell Signaling) or rabbit anti-phospho-STAT6 (Cell Signaling) plus mouse anti-ꞵ-actin (Cell Signaling). Blots were imaged using an Odyssey infra-red imaging system (LI-COR), total signal intensity quantified in Image Studio v5.2 (LI-COR), and the ratio of anti-phospho-STAT6 signal to anti-STAT6 signal calculated. Three biological replicates were carried out. Differences in the phospho-STAT6/STAT6 ratios between strains were tested by two-sided *t-*test with Bonferroni correction.

### Plaque assay

100 parasites were inoculated onto a T25 flask of confluent HFFs and left undisturbed for 10 days. The flasks were stained with 0.5% w/v crystal violet (Sigma), 0.9% w/v ammonium oxalate (Sigma), 20% v/v methanol in distilled water for 15 min then washed with DPBS. To measure knockout efficiency, the number of plaques in flasks supplemented with 5 μM 5-fluorodeoxyuridine were counted relative to untreated flasks in three biological replicates. Differences in percentage knockout efficiency were tested by two-sided *t-*test with Bonferroni adjustment. To measure plaque size, images were taken using a GelDoc Go System (Bio-Rad) and plaque area quantified using ImageJ v1.53c^75^. Differences in plaque sizes between strains was tested by two-sided *t-*test with Bonferroni adjustment, using all measured plaques from one biological replicate.

### Immunofluorescence assays

#### Localisation of GRA59 and SOS1

Confluent HFFs in 8-well chamber slides (Ibidi) were infected with *T. gondii* strains for 24 h then washed with PBS and fixed with 4% w/v formaldehyde (Sigma) for 15 min. The cells were permeabilised with 0.2% Triton X-100 (Sigma) for 15 min and blocked with 2% w/v BSA (Sigma) for 1 h. The cells were stained with 1:500 rat anti-HA (Sigma) followed by 1:1000 goat anti-rat Alexa 594 (Thermo). The cells were then stained with 1:200 mouse anti-*T. gondii* (Santa Cruz) or 1:1000 rabbit anti-*T. gondii* (Abcam) plus 1:1000 mouse anti-GRA1 (Dubremetz lab) or 1:1000 mouse anti-GRA2 (Dubremetz lab) or 1:1000 rabbit anti-GRA3 (Dubremetz lab) or 1:500 mouse anti-ROP1 (Abnova). Finally, the cells were strained with 5 µg/mL DAPI plus 1:1000 goat anti-mouse Alexa 647 (Thermo) and 1:1000 goat anti-rabbit Alexa 488 (Thermo), where the mouse anti-*T. gondii* primary antibody was used, or with 1:1000 goat anti-mouse Alexa 488 (Thermo) and 1:1000 goat anti-rabbit Alexa 647 (Thermo) where the rabbit anti-*T. gondii* primary antibody was used. All antibody incubations were carried out for 1 h at room temperature. Images were acquired on a Nikon Ti-E inverted widefield fluorescence microscope with a Nikon CFI APO TIRF 100x/1.49 objective and Hamamatsu C11440 ORCA Flash 4.0 camera running NIS Elements (Nikon).

#### c-Myc

HFFs were serum starved for 24 h (0.1% FBS), then infected for 24 h, fixed, permeabilised, and blocked as above. The cells were stained with 1:800 rabbit anti-cMyc (Cell Signaling) and 1:200 mouse anti-*T. gondii* (Santa Cruz) for 1 h, followed by 1:1000 goat anti-rabbit Alexa 488 (Thermo), 1:1000 goat anti-mouse Alexa 647 (Thermo), and 5 µg/mL DAPI for 1 h. For each sample, a 3×3 tiled image was acquired as above using a Nikon Plan APO 40x/0.95 objective. cMyc fluorescence intensity was quantified in ImageJ. DAPI signal was used to generate a mask for each host nucleus and the median cMyc intensity in each nucleus of infected cells was measured with the median background (non-nucleus) intensity subtracted. The median nuclear signal per sample was taken as representative of a replicate, and normalised by scaling to RHΔKU80 = 1 AU. Five biological replicates were carried out using HFFs prepared independently and infected on different days. Differences between strains were tested by two-sided Wilcoxon rank-sum test with Bonferroni correction.

#### EZH2

HFFs were infected for 24 h, fixed, permeabilised, and blocked as above. The cells were stained with 1:200 mouse anti-EZH2 (BD) and 1:1000 rabbit anti-*T. gondii* (Abcam) overnight at 4 °C, followed by 1:1000 goat anti-mouse Alexa 488 (Thermo), 1:1000 goat anti-rabbit Alexa 647 (Thermo), and 5 µg/mL DAPI for 1 h. Images were acquired and nuclear EZH2 fluorescence in infected cells was quantified as above for cMyc. Five biological replicates were carried out using HFFs prepared independently and infected on different days. Differences between strains were tested by two-sided Wilcoxon rank-sum test with Bonferroni correction.

#### Phospho-p38

HFFs were infected for 24 h, fixed, permeabilised, and blocked as above. The cells were stained with 1:800 rabbit anti-phospho-p38 (Cell Signaling) and 1:200 mouse anti-*T. gondii* (Santa Cruz) overnight at 4 °C, followed by 1:1000 goat anti-rabbit Alexa 488 (Thermo), 1:1000 goat anti-mouse Alexa 647 (Thermo), and 5 µg/mL DAPI for 1 h. Images were acquired and nuclear phospho-p38 fluorescence in infected cells was quantified as above for cMyc. Five biological replicates were carried out using HFFs prepared independently and infected on different days. Differences between strains were tested by two-sided Wilcoxon rank-sum test with Bonferroni correction.

#### IRF1

HFFs were infected and 50 ng/mL (∼100 U/mL) recombinant human IFNγ added 1 hpi. The cells were fixed 24 hpi, permeabilised, and blocked as above. The cells were stained with 1:200 rabbit anti-IRF1 (Cell Signaling) and 1:200 mouse anti-*T. gondii* (Santa Cruz) overnight at 4 °C, followed by 1:1000 goat anti-rabbit Alexa 488 (Thermo), 1:1000 goat anti-mouse Alexa 647 (Thermo), and 5 µg/mL DAPI for 1 h. Images were acquired and nuclear IRF1 fluorescence in infected cells was quantified as above for cMyc, normalising to uninfected = 1 AU. Five biological replicates were carried out using HFFs prepared independently and infected on different days. Differences between strains were tested by two-sided Wilcoxon rank-sum test with Bonferroni correction.

#### Phospho-STAT6

HFFs were infected for either 1 h or for 24 h, with remaining extracellular parasites washed off at 1 hpi. The cells were fixed with 100% methanol for 15 min and blocked with 2% w/v BSA for 1 h. The cells were stained with 1:200 rabbit anti-phospho-STAT6 (Cell Signaling) and either 1:200 mouse anti-*T. gondii* (Santa Cruz) or 1:10,000 mouse anti-SAG1 (Laboratory of John Boothroyd) overnight at 4 °C, followed by 1:1000 goat anti-rabbit Alexa 488 (Thermo), 1:1000 goat anti-mouse Alexa 647 (Thermo), and 5 µg/mL DAPI for 1 h. Images were acquired and nuclear phospho-STAT6 fluorescence in infected cells was quantified as above for cMyc. Five biological replicates were carried out for each timepoint using HFFs prepared independently and infected on different days. Differences between strains were tested by two-sided Wilcoxon rank-sum test with Bonferroni correction.

#### GRA16-HA/IST-HA export

HFFs were infected for 24 h, fixed with formaldehyde, permeabilised, and blocked as above. The cells were stained with 1:500 rat anti-HA (Sigma), followed by 1:1000 goat anti-rat Alexa 594 (Thermo), followed by 1:1000 rabbit anti-*T. gondii* (Abcam), followed by 1:1000 goat anti-rabbit Alexa 647 (Thermo). For each sample, a 3×3 tiled image was acquired as above using a Nikon Plan APO 40x/0.95 objective. The proportion of exported GRA16-HA/IST-HA was quantified using ImageJ. DAPI signal was used to generate a mask for each host nucleus and anti-*T. gondii* signal was used to generate a mask for each vacuole. The median anti-HA signal in the background (non-nucleus and non-vacuole) was subtracted across the whole image, and the total anti-HA signal in the host nuclei and vacuoles was measured. Export ratio was calculated as the amount of GRA16-HA/IST-HA immunofluorescence signal in the host nuclei compared to that remaining in the vacuoles. To determine the percentage of vacuoles with IST-HA accumulations, the images were blinded and the vacuoles manually scored for the presence/absence of IST-HA accumulations. Five biological replicates were carried out using HFFs prepared independently and infected on different days. Differences between strains were tested by unpaired, two-sided *t-*test with Bonferroni correction.

### Arginase activity assay

10^6^ BMDMs per sample were infected with *T. gondii* parasites at a MOI of 1 for 36 h. Arginase activity in the cell lysate was measured using a colorimetric assay (Abcam). Two biological replicates were carried out with BMDMs prepared and infected on different days, with triplicate infections for each strain in each biological replicate. Differences between strains were tested by two-sided Wilcoxon rank-sum test with Bonferroni correction.

### Identification of protein homologues

The AlphaFold structural prediction for SOS1 (TGGT1_222100) was accessed from the AlphaFold/EMBL-EBI database version 2022-11-01^76, 77^. Homologues of SOS1 were identified using HMMER^78^ with UniProtKB reference proteomes with an E-value cut-off of 0.01. No significant hits were found outside of Apicomplexa. Regions of >40% similarity (BLOSUM62) were identified by BLAST via VEuPathDB^31^. A phylogenetic tree of select homologues was generated by Clustal Omega^79^.

### Cyst formation assay

Cyst formation assays were carried out as previously described^60^. Briefly, primary neurons were cultured in 96-well, clear-bottom, black-walled plates and infected with indicated *T. gondii* strains at a MOI of 0.1. At 1, 2, and 3 days post-infection, the cells were fixed with 4% w/v formaldehyde for 15 min, then simultaneously permeabilised and blocked with 0.1% v/v Triton X-100 plus 3% w/v goat serum.

The cells were stained with 1:10,000 rabbit anti-*T. Gondi* (Thermo) and 1:500 biotinylated *Dolichos bisflorus* agglutinin (DBA) (Vector Labs) for 1 h, followed by 1:500 streptavidin Alexa 647 (Thermo), 1:500 goat anti-rabbit Alexa 488 (Thermo) and DAPI (Thermo) for h. Wells were covered with 100 μLs of 1:50 Fluoromount-G (Thermo) for imaging. Entire wells were imaged at 20x magnification (69 fields of view per well) using an Operetta high content imaging system (PerkinElmer). Image analysis was carried out using Harmony (PerkinElmer). *T. gondii* vacuoles were identified by anti-*T. gondii* staining. Only vacuoles containing two or more parasites (defined by a minimum vacuole length of 8 µm and width of 4 µm) were considered for further analysis. The percentage of DBA-positive vacuoles was determined in each well, and the mean percentage across replicate wells for each strain taken to represent each biological replicate. Three biological replicates were carried out. Differences between strains were determined by two-sided *t*-test with Bonferroni correction.

## RESOURCE AVAILABILITY

### Lead contact

Further information and requests for reagents may be directed to and will be fulfilled by the lead contact, Moritz Treeck.

### Materials availability

All plasmids and *T. gondii* cell lines generated in this study are available upon request.

### Data and code availability

Single cell RNA-sequencing data (demultiplexed FASTQ files and Cell Ranger count matrices) have been deposited at the Gene Expression Omnibus with the accession number GSE229505 and will be made publicly available prior to final publication.

All original code and supporting files used to analyse the single-cell RNA-sequencing data in this paper are available on GitHub at https://github.com/simonwbutterworth/Dual-Perturb-Seq-2023/ and will be made publicly available prior to final publication

Any additional information required to reanalyse the data reported in this paper is available upon request.

## ACKNOWLEDGEMENTS

We thank the Cell Services, High Throughput Screening, Flow Cytometry, and Advanced Sequencing Facility Science Technology Platforms at the Francis Crick Institute for support. We thank the laboratory of Dominique Soldati for providing 49c. We thank Manfred Claassen and Revant Gupta for discussion and advice on data analysis. We thank VEuPathDB^31^ for providing access to genomic and other large-scale datasets for *T. gondii*.

This work was supported by an award to MT from the Wellcome Trust (223192/Z/21/Z), by funding to MT from the Francis Crick Institute, which receives its core funding from Cancer Research UK (CC2132), the UK

Medical Research Council (CC2132), and the Wellcome Trust (CC2132), by funding to AK from the College of Medicine, University of Arizona, and by funding to AK from the BIO5 Institute, University of Arizona. FT is funded by the Deutsche Forschungsgemeinschaft (TO 1349/1-1). The Science Technology Platforms at the Francis Crick Institute receive funding from Cancer Research UK (CC0199), The UK Medical Research Council (CC0199) and the Wellcome Trust (CC0199).

For the purpose of Open Access, the authors have applied a CC BY public copyright licence to any Author Accepted Manuscript version arising from this submission.

## AUTHOR CONTRIBUTIONS

Conceptualisation: SB, MT. Methodology: SB, AE. Investigation: SB, SC, KT, FT, EL. Formal analysis: SB, KK. Visualisation: SB. Writing - original draft: SB, MT. Writing - review and editing: all authors. Supervision: RG, AK, MT. Funding acquisition: AK, MT.

## DECLARATION OF INTERESTS

The authors declare no competing interests.

## SUPPLEMENTARY FIGURES

**Figure S1.**
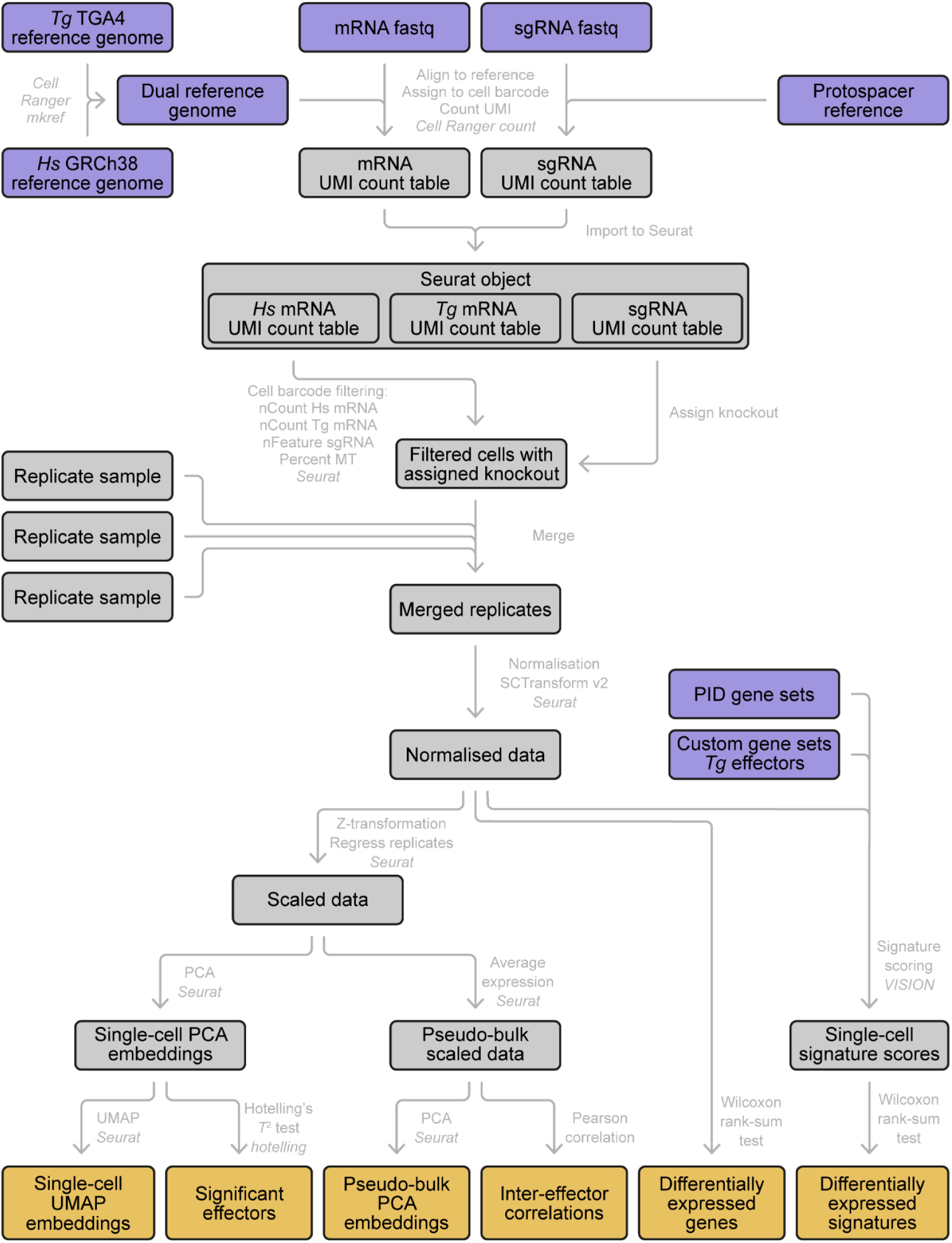
Schematic of dual perturb-seq data processing and analysis.

**Figure S2.**
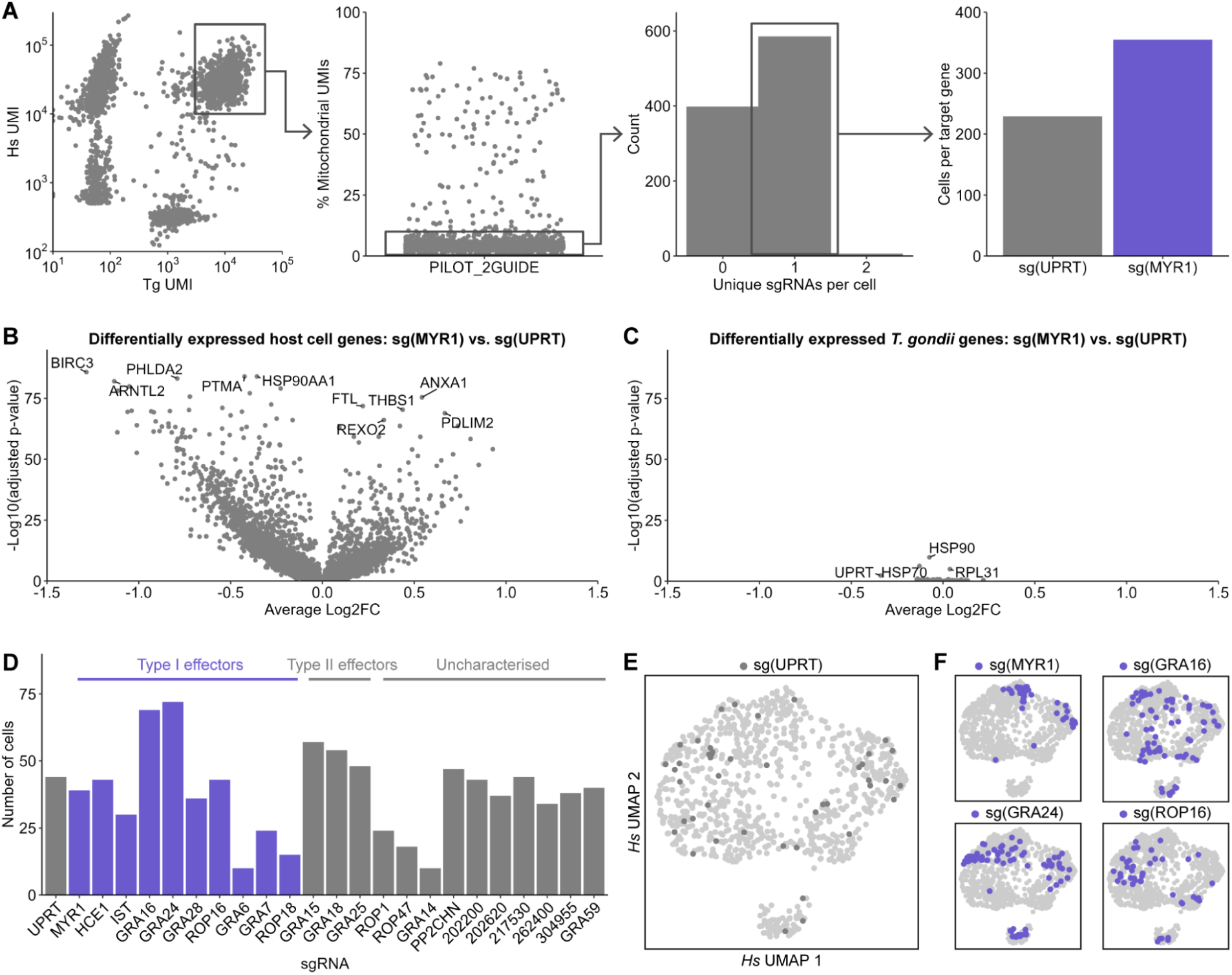
Filtering and assignment of sgRNA identity to dual perturb-seq transcriptomes. **A.** Typical cell barcode filtering strategy for dual perturb-seq data. **B.** Differentially expressed host cell genes for sg(MYR1)-expressing cells compared to sg(UPRT)-expressing cells (two-sided Wilcoxon rank-sum test with Benjamini-Hochberg adjustment). See also **Supplementary Data 1A**. **C.** Differentially expressed *T. gondii* genes for sg(MYR1)-expressing cells compared to sg(UPRT)-expressing cells (two-sided Wilcoxon rank-sum test with Benjamini-Hochberg adjustment). See also **Supplementary Data 1B**. **D.** Number of single cell transcriptomes recovered for each target gene in 24-sgRNA pilot experiment. **E.** Distribution of sg(UPRT)-expressing cells in UMAP of host cell gene expression in 24-sgRNA pilot experiment. **F.** Distribution of cells expressing sgRNAs targeting select effectors in UMAP of host cell gene expression in 24-sgRNA pilot experiment.

**Figure S3.**
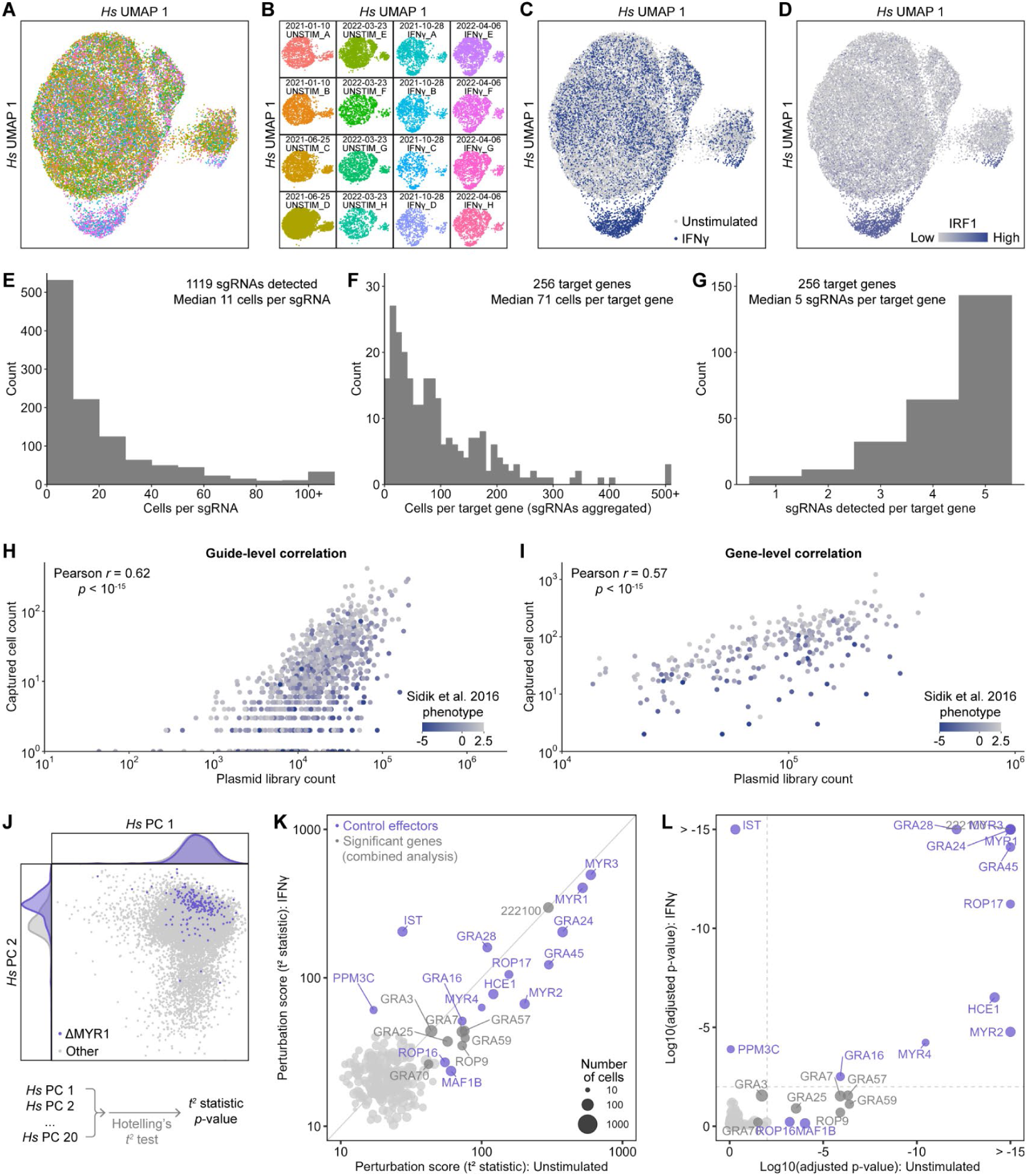
Quality control analysis of dual perturb-seq screen. **A.** UMAP of single cell transcriptomes based on host cell gene expression coloured by sample. **B.** UMAP of single cell transcriptomes split by sample. **C.** UMAP of single cell transcriptomes coloured by condition (unstimulated or stimulated with IFNγ). **D.** Expression of the interferon-stimulated gene IRF1. **E.** Histogram of the number of single cell transcriptomes expressing each sgRNA. See also **Supplementary Data 3**. **F.** Histogram of the number of single cell transcriptomes for each target gene. **G.** Histogram of the number of sgRNAs detected for each target gene. **H.** Correlation between the number of read counts in bulk sequencing data of perturb-seq plasmid library and the number of single cell transcriptomes for each sgRNA. See also **Supplementary Data 3**. **I.** Correlation between the number of read counts in bulk sequencing data of perturb-seq plasmid library and the number of single cell transcriptomes summed for each target gene. **J.** Illustration of Hotelling’s *t*^2^-test on single cell PCA embeddings. **K.** Correlation between perturbation scores (Hotelling’s *t*^2^-test statistic) in unstimulated and IFNγ-stimulated samples. See also **Supplementary Data 4**. **L.** Correlation between Hotelling’s *t*^2^-test p-values in unstimulated and IFNγ-stimulated samples. See also **Supplementary Data 4**.

**Figure S4.**
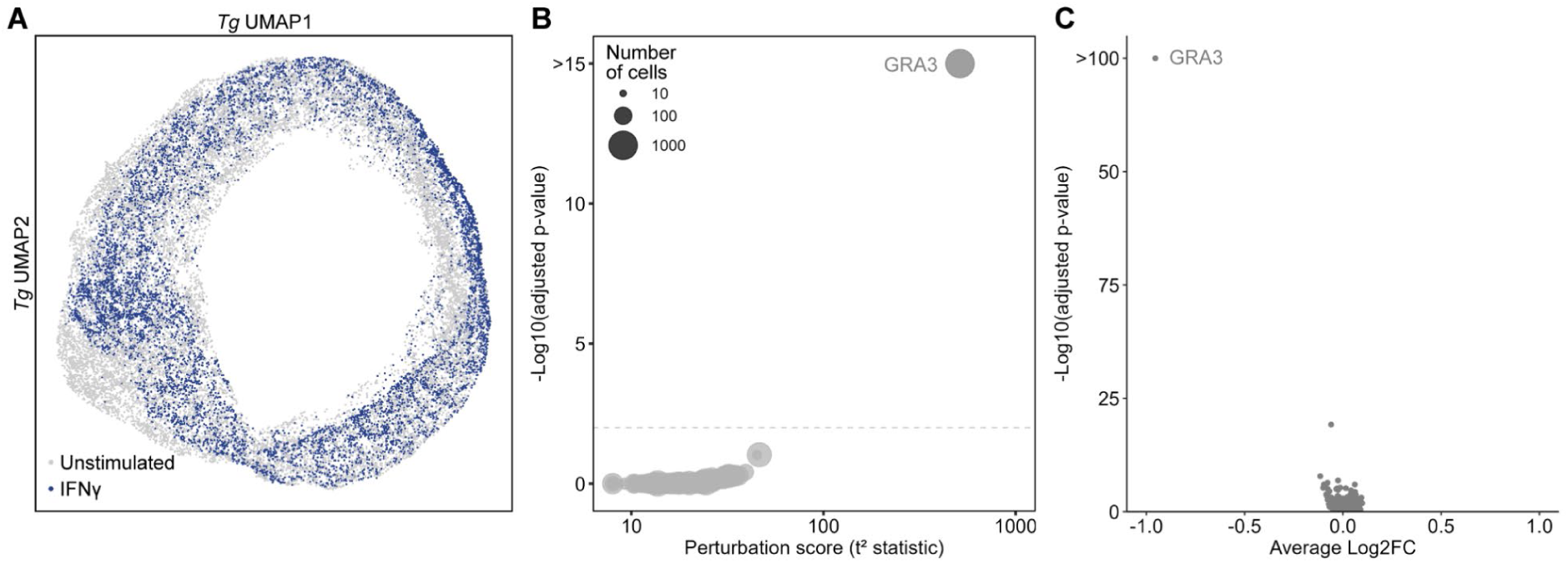
Perturbation of the *T. gondii* transcriptome by effector proteins. **A.** UMAP of single cell transcriptomes based on *T. gondii* gene expression with cells coloured by condition (unstimulated or stimulated with IFNγ). **B.** Perturbation of *T. gondii* transcriptome by effectors, measured by Hotelling’s *t*^2^ test on PCA embeddings of single cell transcriptomes with Benjamini Hochberg adjustment. See also **Supplementary Data 5.** **C.** Differentially expressed *T. gondii* genes for sg(GRA3)-expressing cells compared to all other cells (two-sided Wilcoxon rank-sum test with Benjamini-Hochberg adjustment). See also **Supplementary Data 6**.

**Figure S5.**
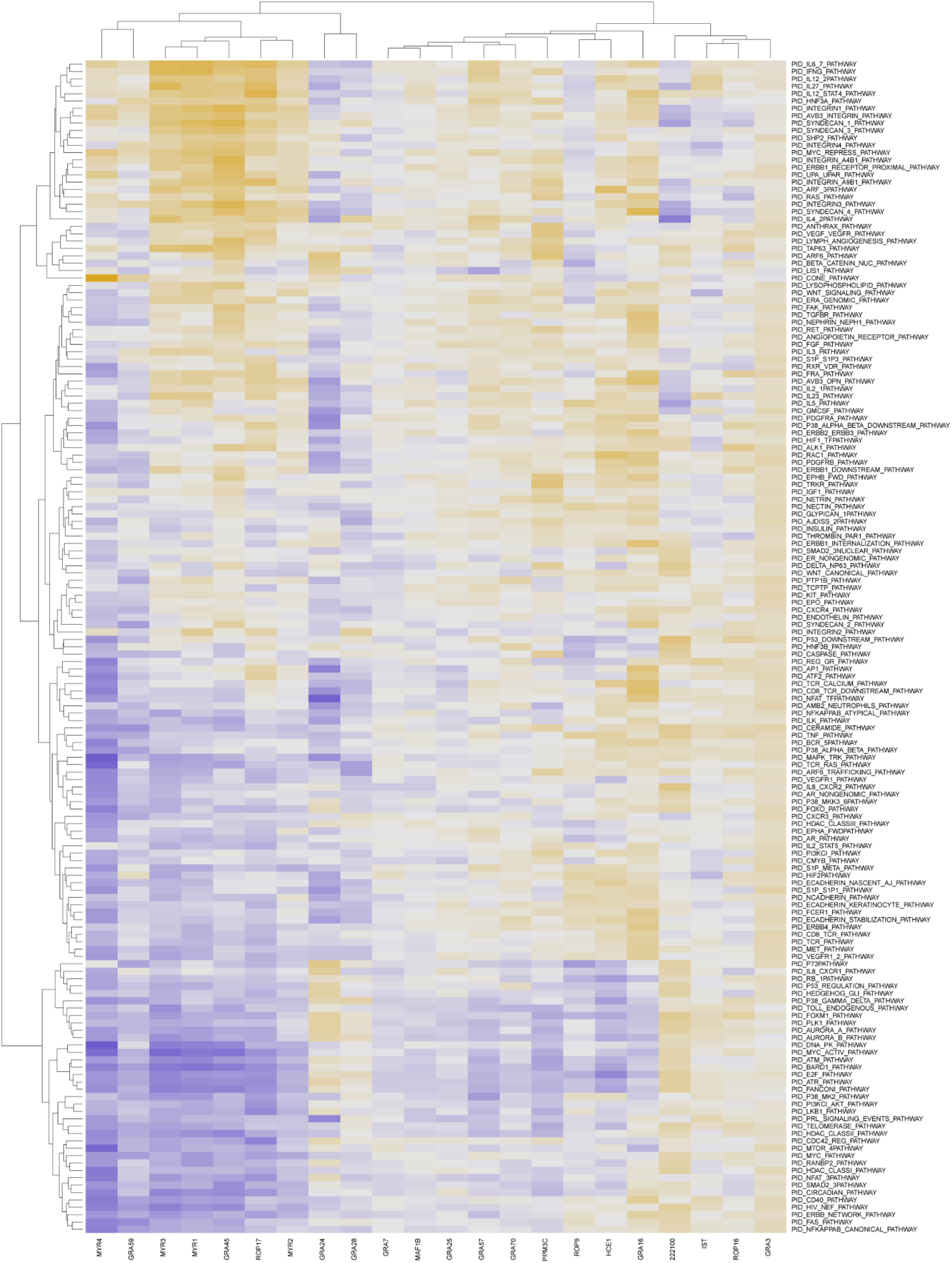
Pathway Interaction Database gene sets that are significantly differentially regulated by *T. gondii* effector proteins. Average VISION signature scores of Pathway Interaction Database gene sets that are significantly differentially regulated by at least one significant effector (p < 0.01, two-sided Wilcoxon rank-sum test with Benjamini-Hochberg adjustment). See also **Supplementary Data 7**.

**Figure S6.**
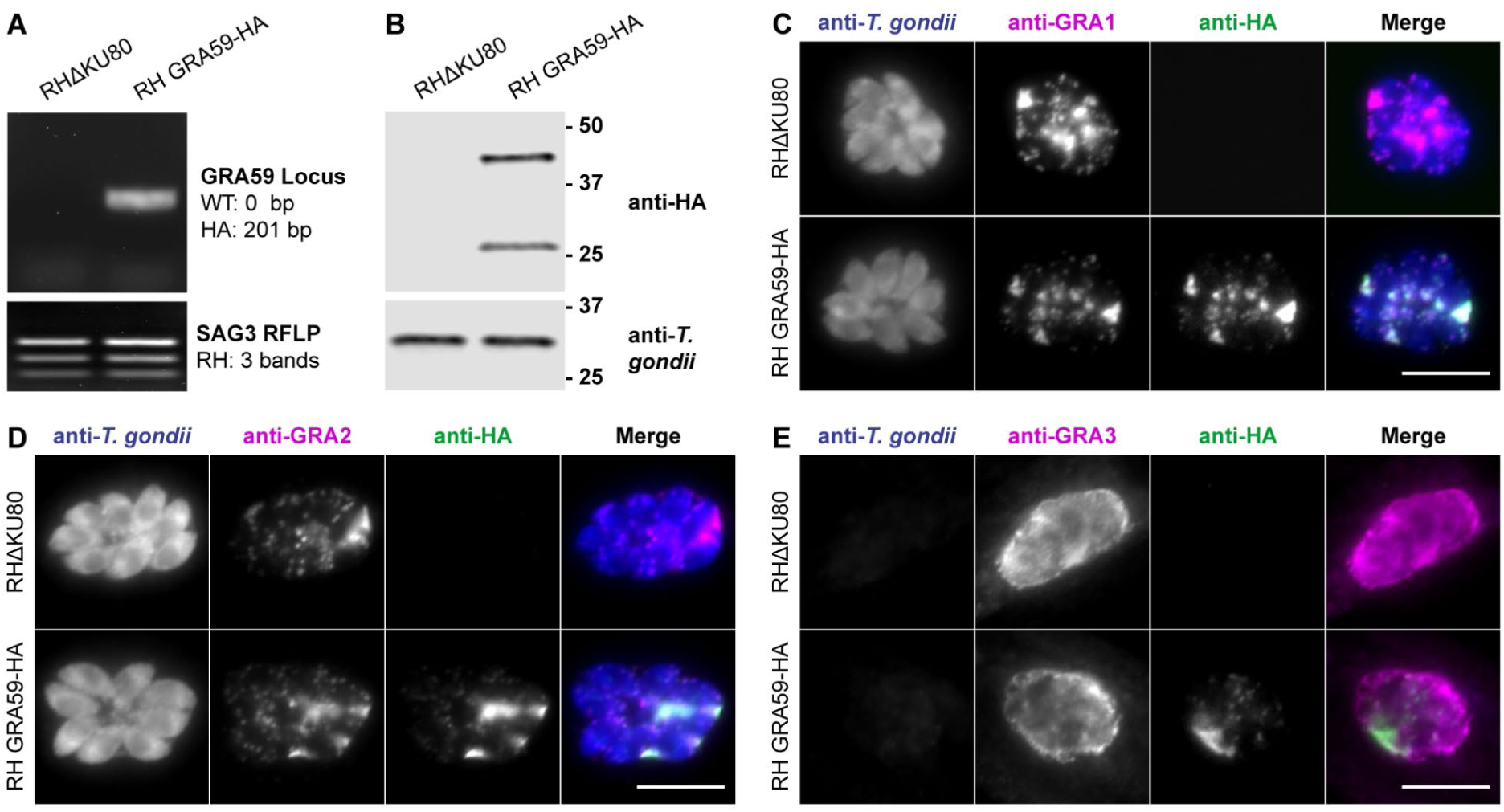
C-terminal epitope tagging of GRA59. **A.** Verification of HA tagging by diagnostic PCR. **B.** Verification of GRA59-HA expression by Western blot. **C.** Co-localisation of GRA59-HA with GRA1 by immunofluorescence assay. Scale bar = 10 µm. **D.** Co-localisation of GRA59-HA with GRA2 by immunofluorescence assay. Scale bar = 10 µm. **E.** Co-localisation of GRA59-HA with GRA3 by immunofluorescence assay in cells permeabilised with 0.1% saponin for 15 min. Scale bar = 10 µm.

**Figure S7.**
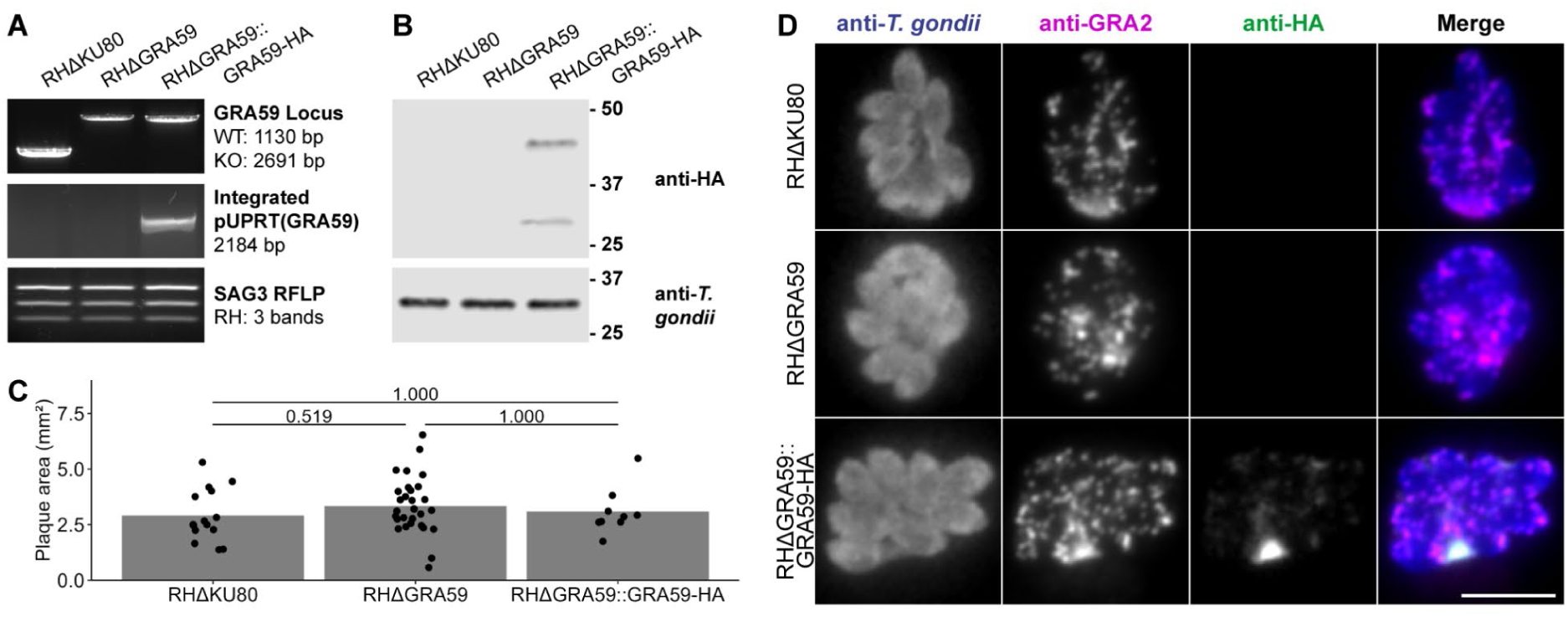
Knockout and complementation of GRA59. **A.** Verification of GRA59 knockout and complementation by diagnostic PCR. **B.** Verification of GRA59-HA expression by Western blot. **C.** Plaque assay for RHΔKU80, RHΔGRA59, and RHΔGRA59::GRA59-HA. One biological replicate; points represent individually measured plaques. Differences tested by two-sided *t*-test with Bonferroni correction. **D.** Verification of GRA59-HA expression and localisation by immunofluorescence assay. Scale bar = 10 µm.

**Figure S8.**
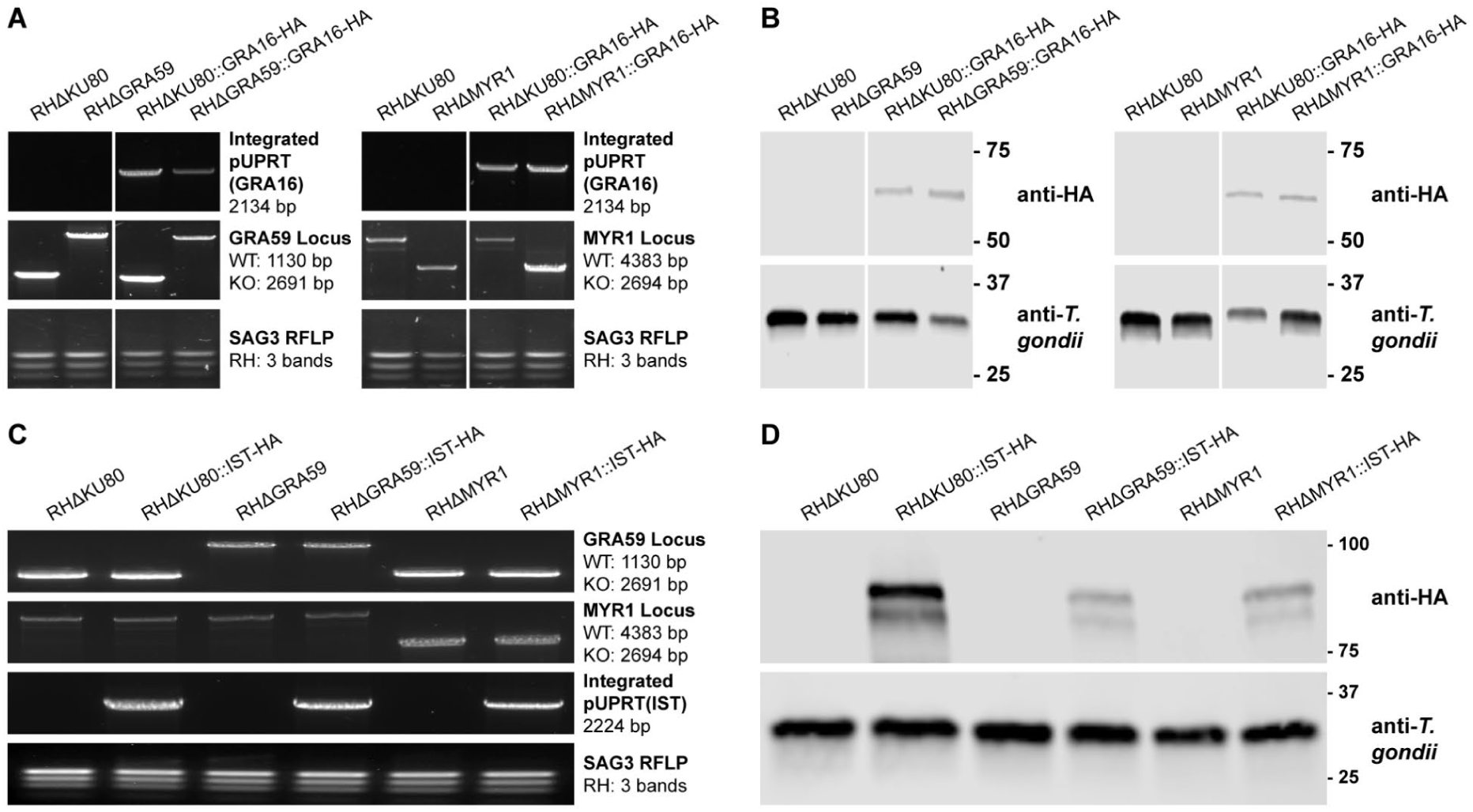
Introduction of GRA16-HA to GRA59 and MYR1 knockout cell lines. **A.** Verification of pUPRT(GRA16-HA) integration by diagnostic PCR. **B.** Verification of GRA16-HA expression by Western blot. **C.** Verification of pUPRT(IST-HA) integration by diagnostic PCR. **D.** Verification of IST-HA expression by Western blot.

**Figure S9.**
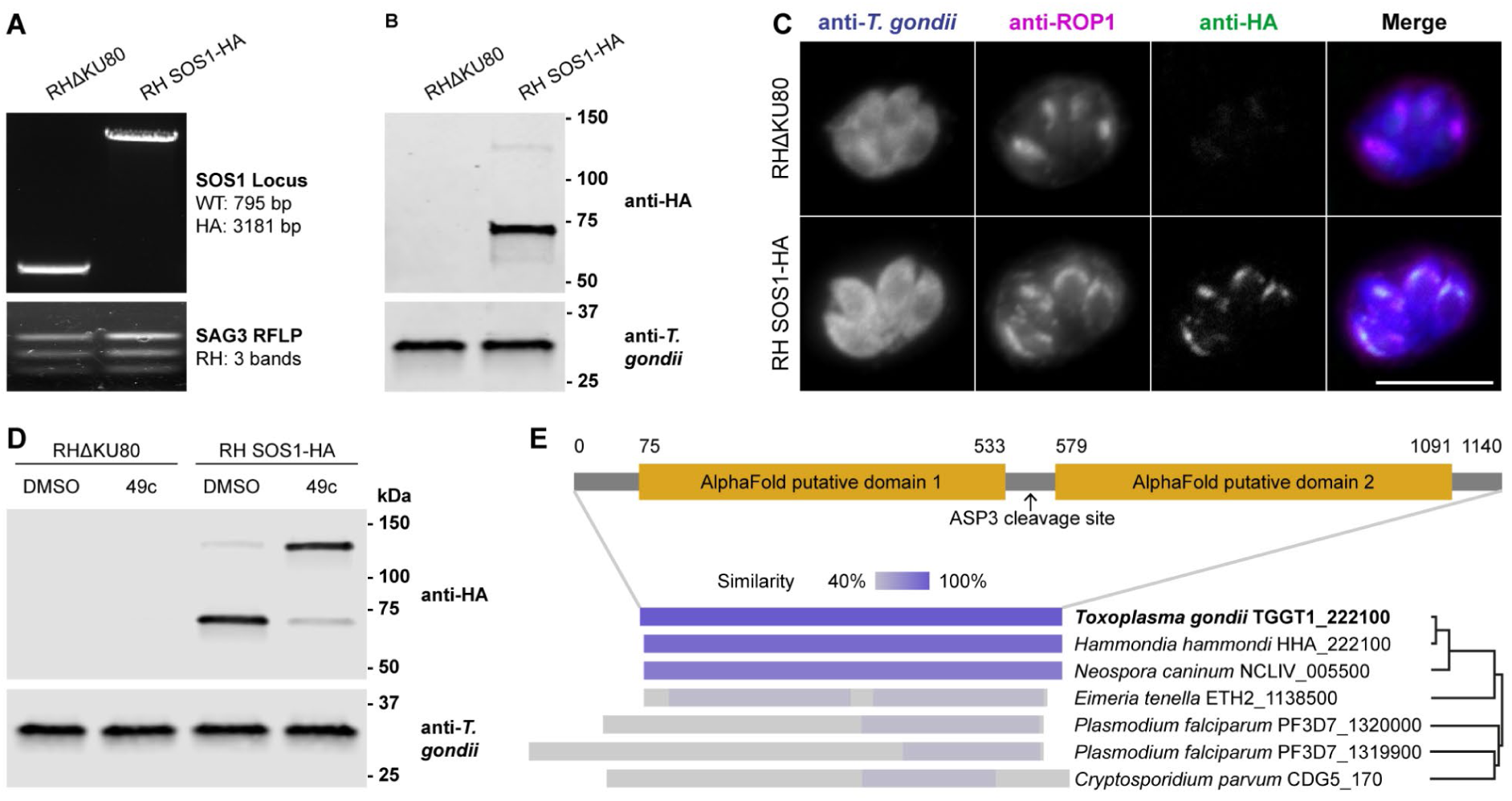
C-terminal epitope tagging of SOS1. **A.** Verification of HA tagging by diagnostic PCR. **B.** Verification of SOS1-HA expression by Western blot. **C.** Co-localisation of SOS1-HA with ROP1 by immunofluorescence assay. Scale bar = 10 µm. **D.** Treatment of parasites with the ASP3 inhibitor 49c reduces processing of SOS1. 10 µM 49c was added at 1 hpi and parasites were harvested by syringe-lysis at 48 hpi. **E.** Putative structure of SOS1 and alignment to homologues detected in Apicomplexa.

**Figure S10.**
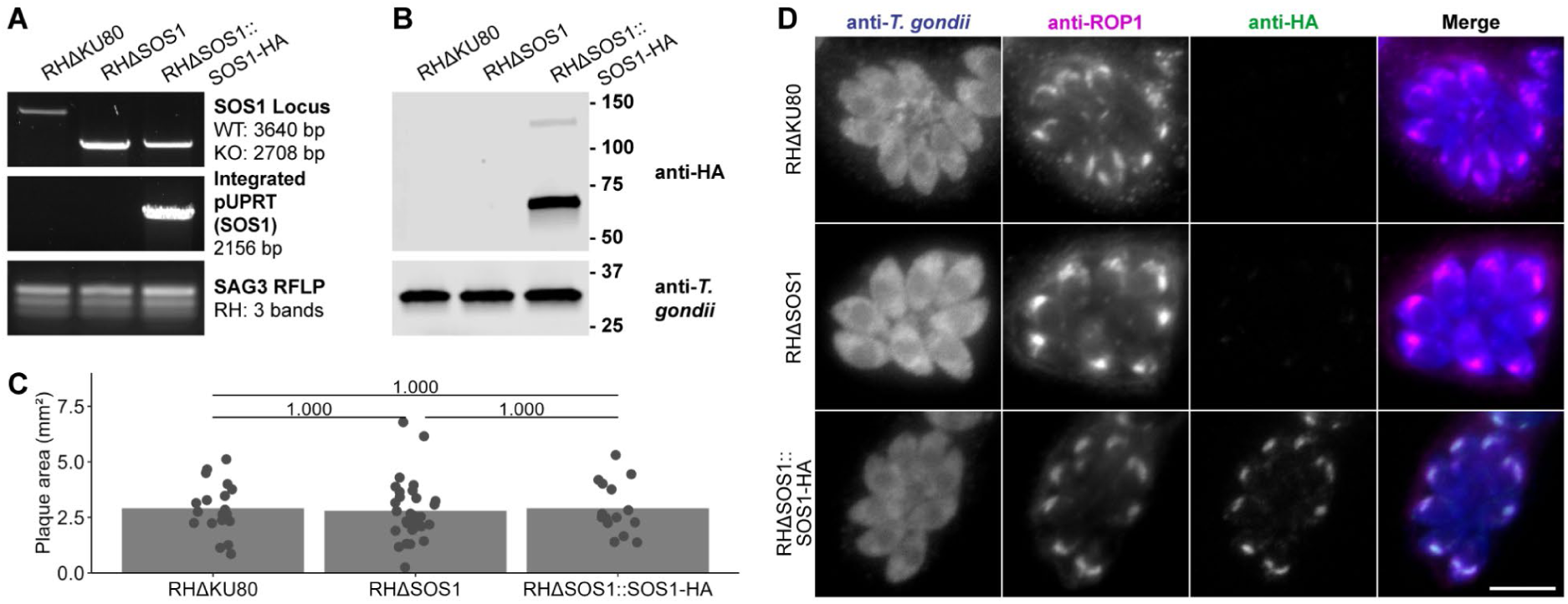
Knockout and complementation of SOS1 in *T. gondii* RH. **A.** Verification of SOS1 knockout and complementation by diagnostic PCR. **B.** Verification of SOS1-HA expression by Western blot. **C.** Plaque assay for RHΔKU80, RHΔSOS1, and RHΔSOS1::SOS1-HA. One biological replicate; points represent individually measured plaques. Differences tested by two-sided *t*-test with Bonferroni correction. **D.** Verification of SOS1-HA expression by immunofluorescence assay. Scale bar = 10 µm.

**Figure S11.**
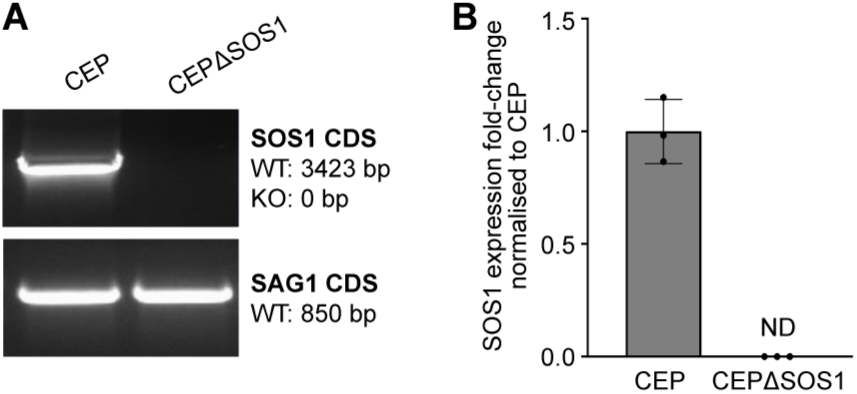
Knockout and complementation of SOS1 in *T. gondii* CEP. **A.** Verification of SOS1 knockout by diagnostic PCR. **B.** qPCR quantification of SOS1 mRNA expression relative to *Tg*Actin.

## SUPPLEMENTARY DATA FILES

**Supplementary Data 1.**

**A.** Differentially expressed host cell genes between sg(MYR1)-and sg(UPRT)-expressing cells.

**B.** Differentially expressed T. gondii genes between sg(MYR1)-and sg(UPRT)-expressing cells.

**Supplementary Data 2. Significant differentially expressed genes identified in 24-guide pilot experiment.**

**Supplementary Data 3. Gene targets and protospacer sequences used in dual perturb-seq screen.**

**Supplementary Data 4. Hotelling’s *t*^2^-test results for host cell gene expression.**

**A.** Results for combined data (unstimulated and IFNγ-stimulated).

**B.** Results for unstimulated sample data.

**C.** Results for IFNγ-stimulated sample data.

**Supplementary Data 5. Hotelling’s *t*^2^-test results for *T. gondii* gene expression.**

**Supplementary Data 6. Differentially expressed *T. gondii* genes for sg(GRA3)-expressing cells.**

**Supplementary Data 7. Differentially expressed PID gene sets.**

**Supplementary Data 8. DNA primer sequences used in this work.**

**Supplementary Data 9. Plasmids generated in this work.**

## Notes

### Competing Interest Statement

The authors have declared no competing interest.

